# Matched Plasma and Tissue Proteomics Reveal the Selective Nature of Circulating Biomarkers of Human Aging

**DOI:** 10.64898/2026.10.19.759783

**Authors:** Qiongrong Huang, Mark A. Watson, John C. Earls, Noa Rappaport, Heather L. Gregory, John Stone, Maryellen G. Pavone, Jamie N. Justice, Bumsoo Ahn, Ryan P. Lamers, Annalise Schweickart, Christopher C. Benz, Francesca E. Duncan, Birgit Schilling, Nathan D. Price

**Author notes:** Corresponding authors: Birgit Schilling and Nathan D. Price. Equal contribution (Qiongrong Huang, Mark A. Watson).

## Abstract

Aging is systemic, yet how circulating proteins reflect molecular aging within solid tissues remains unclear. We profiled 5,415 proteins using Olink Explore HT in 178 adults aged 20-87 years, including matched plasma-muscle (n = 56) and plasma-ovary (n = 24) pairs. Plasma-tissue concordance was limited, with 50 muscle and 17 ovarian proteins significantly correlated, indicating that plasma selectively reflects tissue protein abundance. Age-correlated proteins and pathways differed markedly across sex and tissue strata, Including immune/inflammatory signatures in female plasma, vascular and extracellular-matrix pathways in male plasma, and collagen remodeling in the ovary. Expression Variation Analysis further revealed predominantly context-specific age-related changes in protein coordination, including FAS/FADD/TRAIL death-receptor networks in plasma and a TP53-CDKN1A senescence-associated axis in muscle. Exploratory multivariate models identified additional plasma associations with subsets of tissue proteins, including tissue-restricted muscle proteins. Together, these findings define plasma as a selective, context-dependent window into human tissue aging.

## Main

Aging is fundamentally a progressive loss of physiological function and resilience, arising from cumulative molecular and cellular damage and culminating in organ dysfunction and increased disease risk^1–3^. Here, we consider resilience as the capacity of biological systems to maintain or restore homeostasis through coordinated responses to physiological challenges. While chronological age provides a crude measure of aging, biological aging occurs heterogeneously across tissues and between individuals^4,5^. However, directly sampling most organs longitudinally in healthy individuals is impractical and often invasive, motivating the development of minimally invasive system-level plasma biomarkers capable of tracking biological aging across an individual. Most human molecular studies of aging focus on average shifts in individual molecules^6–8^. However, organismal resilience depends on the coordinated activity of protein programs that regulate immunity, metabolism, tissue maintenance and repair. With age, biological systems become increasingly heterogeneous and regulatory processes more stochastic, suggesting that aging may manifest not only as changes in mean protein abundance but also as altered coordination among proteins. Quantifying both abundance and coordination can therefore provide complementary views of biological aging.

Blood plasma integrates secreted and leaked proteins from many tissues^9,10^, enabling minimally invasive monitoring. Large-scale plasma proteomics has revealed age-correlated trajectories across the lifespan^11^ and proteomic clocks that predict mortality and age-related disease^12^. Nevertheless, the extent to which circulating proteins faithfully reflect molecular aging within specific tissues remains unclear^9,13^. Most approaches infer tissue states from plasma without matched tissues measurements^14^, while studies employing paired plasma and tissue samples have largely focused on disease settings, particularly in cancer^10^, where tissue perturbations are pronounced and readily detectable in blood. Matched plasma-tissue studies in healthy individuals remain uncommon because tissue biopsies are rarely obtained outside clinical indications, and existing multi-tissue atlases typically provide static snapshots^9^, limiting our understanding of how physiological tissue aging is represented in the circulating proteome. A central unanswered question is therefore not simply which proteins change with age in plasma, but whether circulating proteomic aging reflects the molecular remodeling occurring within individual organs. We focused on muscle and ovary because they represent distinct, clinically relevant modes of functional aging. Muscle is a continuously active and highly plastic organ system that underpins mobility and whole-body metabolic homeostasis and is central to frailty risk^15–17^. In contrast, the ovary is a sex-specific organ that exhibits some of the earliest overt signs of aging and continues to undergo molecular and tissue remodeling beyond the reproductive period^18,19^. Together, these tissues provide complementary contexts for determining when and how organ-specific aging signals become measurable in plasma. Whether age-correlated changes observed in plasma reflect molecular aging within specific tissues, and conversely whether tissue aging leaves any detectable signatures in plasma, remains largely unknown.

Here, we advance our understanding of how circulating proteomic aging reflects organ-specific molecular aging by performing paired human plasma–tissue proteomics in 178 adults spanning 20–87 years. We quantified 5,415 proteins (Olink Explore) in plasma (n=169), muscle (n=57), and ovary (n=30), including matched plasma–muscle and plasma–ovary samples from the same individuals (**Fig. 1 and Table 1**). This design enabled us to determine (i) the extent to which between-individual differences in tissue protein abundance are reflected in plasma; (ii) how sex and organ shape age-correlated proteomic and network remodeling; and (iii) whether multivariate plasma profiles contain information about tissue protein abundance beyond direct matched-protein concordance.

**Fig. 1:**
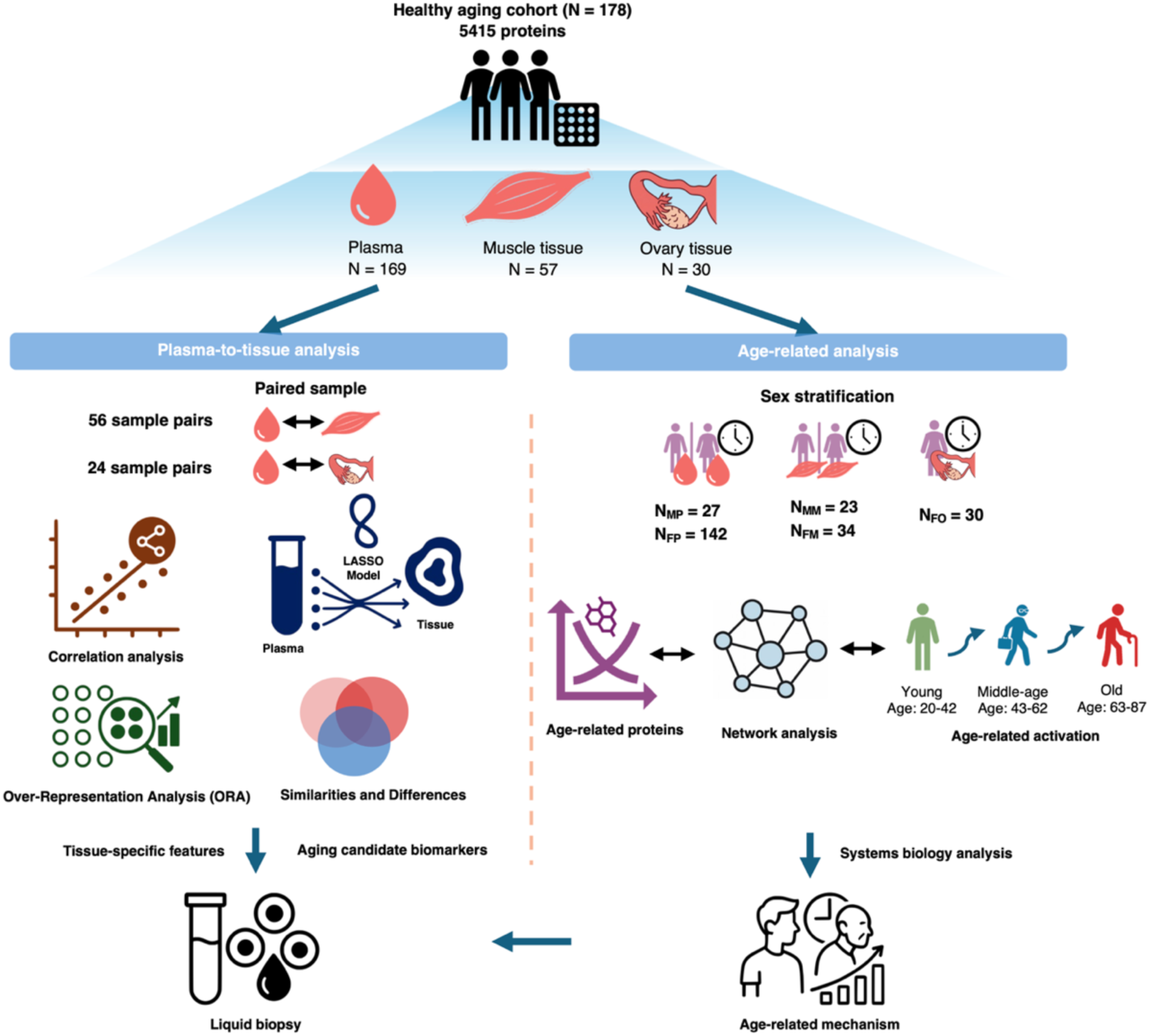
Study overview. Schematic of the study design in 178 adults aging cohort (20–87 years), in which 5,415 proteins were quantified across plasma (N = 169), muscle tissue (N = 57), and ovary tissue (N = 30). Two complementary analytical frameworks were implemented. The plasma–tissue paired analysis used 56 matched muscle–plasma pairs and 24 matched ovary–plasma pairs to quantify cross-compartment concordance (Spearman correlation), apply LASSO regression to predict tissue abundance from plasma, and perform over-representation analysis (ORA) to identify tissue-specific features, thereby informing circulating biomarker discovery for plasma biomarker. The sex- and tissue-stratified age-correlated analysis divided the cohort into male plasma (N = 27), female plasma (N = 142), male muscle (N = 23), female muscle (N = 34), and ovary (N = 30), and further stratified by age group: young (20–42 yr), middle-age (43–62 yr), and old (63–87 yr). This analysis identified age-correlated proteins and interrogated their network architecture using Expression Variation Analysis (EVA), with biological processes inferred via STRING.

**Table 1.**
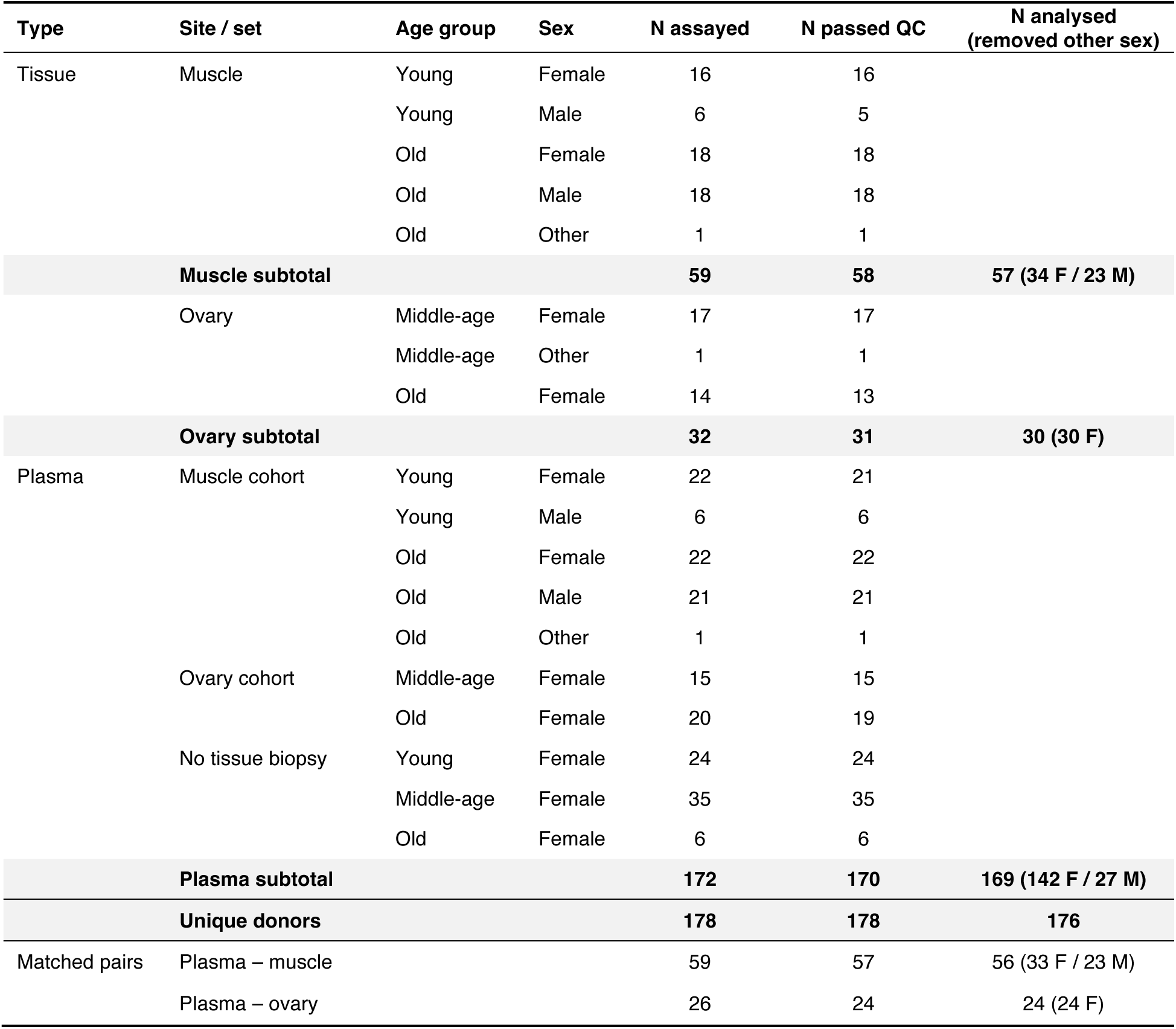
Sample overview and accounting. Breakdown of the study cohort by sample type, collection site, age group (young 20–42, middle-age 43–62, old 63–87 yr) and sex, followed by the matched plasma–tissue sets. N assayed, samples run on Olink Explore HT (control wells excluded); for the matched-pair rows, participants with both compartments collected. N passed QC, samples retained after assay- and sample-level QC and after removal of samples flagged as outliers; for the matched-pair rows, participants retained in both QC-passed matrices. N analysed, the set entering the sex-stratified analyses. Participant totals are unchanged by QC (178 before and after), because each of the four samples removed at QC came from a participant retained in the other compartment. Of the 178 participants, 176 enter the sex- and tissue-stratified analyses (149 female, 27 male); the remaining two are the participants recorded as Sex = Other, which cannot be assigned to a sex stratum. Of the 57 quality-controlled plasma–muscle participant pairs, 56 enter the paired correlation and LASSO analyses; the excluded pair is the participant recorded as Sex = Other.

## Results

### A matched human plasma–tissue proteomic cohort spanning human aging

To investigate how circulating proteins reflect tissue-specific molecular changes during human aging, we assembled a cohort of 178 individuals aged 20–87 years (149 females, 27 males, and 2 individuals of other gender; the latter were excluded from sex-stratified analyses), with non-pathological tissues sampled for proteomic profiling and quantified 5,415 proteins using the Olink Explore platform across plasma (n=169), muscle (n=57), and ovarian tissue (n=30) (**Fig. 1a and Table 1**). All muscle participants and 26 ovary participants also provided plasma, 178 unique participants corresponded to the 172 plasma participants plus the 6 ovary participants from whom no plasma was drawn. Within the plasma set entering sex-stratified analyses (n = 169) the split was 142 female and 27 male; the seven remaining female participants either contributed ovarian tissue only (n = 5) or had their plasma sample removed at QC while their tissue sample was retained (n = 2) (**Table 1**). Matched sampling provided 56 plasma-muscle pairs and 24 plasma-ovary pairs for within-individual comparisons following a quality-control filtering (**Methods**).

We used two complementary analytical frameworks (**Fig. 1a**). First, matched plasma–tissue pairs were analyzed to quantify plasma–tissue concordance and to explore whether multivariate plasma profiles contain information about tissue protein abundance beyond direct concordance. Second, sex- and tissue-stratified analyses were performed to identify age-correlated proteins and characterize age-related network remodeling using Expression Variation Analysis (EVA), a rank-based approach that quantifies changes in protein co-regulation within predefined protein modules. For EVA, samples were grouped by age according to tissue availability. Female plasma samples were divided into young (20-42 years, n = 45), middle-aged (43-62 years, n = 50) and old (63-87 years, n = 47) groups. Male plasma and skeletal muscle were analyzed as young versus old (male plasma: 23-30 years, n = 6 versus 70-87 years, n = 21; female muscle: 20-35 years, n = 16 versus 70-87 years, n = 18; male muscle: 23-30 years, n = 5 versus 70-87 years, n = 18), whereas ovarian tissue, which undergoes age-related functional and molecular remodeling earlier than many other organs, was analyzed as middle-aged (43-62 years, n = 17) versus old (63-87 years, n = 13) (see **Methods** and **Table 1** for details).

Study-site effects in the female plasma arm were adjusted using ComBat while preserving biological variation attributable to age and body mass index (BMI); other strata were each collected at a single site and were not batch-corrected, as confirmed by variance partitioning analysis using the variancePartition R package before and after correction (**Supplementary Fig. 1**). Because ComBat was applied to the female plasma arm only, and because samples were collected across different sites, every sex comparison reported below was repeated using only participants from the single site from which all 27 male participants originated, without batch correction, and is reported alongside the all-site analysis. For the EVA input, ComBat was applied on the log₂ scale; the log₂ transformation alone preserves ranks, whereas ComBat adjustment can alter between-protein ranks.

### Limited plasma-tissue concordance reveals the selective nature of circulating biomarkers of human aging

Following preprocessing and quality control (**Methods**), we calculated within-protein Spearman correlations between matched plasma and tissue, adjusted for age, sex and BMI. These correlations quantify covariation after age adjustment, rather than concordance of age-related changes. Plasma-tissue concordance was generally low, with correlation coefficients centered near zero for both ovary and muscle (**Fig. 2a, b**), indicating that most tissue proteins showed little correspondence with their circulating counterparts. For ovary, plasma-tissue correlations had a median Spearman’s ρ = 0.08 (range, −0.63 to 0.85). Only 17 of 5,415 proteins (0.3%) remained significant after Benjamini-Hochberg correction (adjusted P < 0.05), all exhibiting strong positive correlations (ρ = 0.71-0.85) that occupied the extreme right tail of the distribution (**Fig. 2a, c and Supplementary Table 1**). Muscle showed similarly limited concordance (median ρ = 0.03; range, −0.47 to 0.84), with 50 of 5,415 proteins (0.9%) significant after correction, comprising 48 positive (ρ = 0.46-0.84) and two negative correlations (ρ = −0.46 to −0.47) (**Fig. 2b, d and Supplementary Table 2**). Among the proteins that showed the strongest plasma-tissue correlations were ABO, CD33, IFNGR2 and SIRPA in ovary, and HDGF, CCL24 and HBZ in muscle (**Fig. 2c, d**). These top-ranked proteins were predominantly secreted factors, immune receptors, and circulating signaling molecules, whereas intracellular structural proteins showed little evidence of plasma-tissue concordance. Because measurements near the limit of detection (LOD) may attenuate observed correlations, we plotted ρ against the mean signal above the Olink fixed LOD separately for each compartment (**Supplementary Fig. 2 and Supplementary Tables 1–2**). Detectability explained little of the spread: the Spearman correlation between |ρ| and mean signal above LOD was 0.12 on the plasma side and 0.09 on the ovary side of the ovary contrast, and 0.10 and 0.07 for muscle, while restricting to proteins detected in ≥ 90% of participants raised the median ρ only from 0.079 to 0.143 in ovary and from 0.028 to 0.057 in muscle. Low concordance persisted among well-detected proteins, suggesting that proximity to the assay LOD alone does not explain the near-zero correlations.

**Fig. 2:**
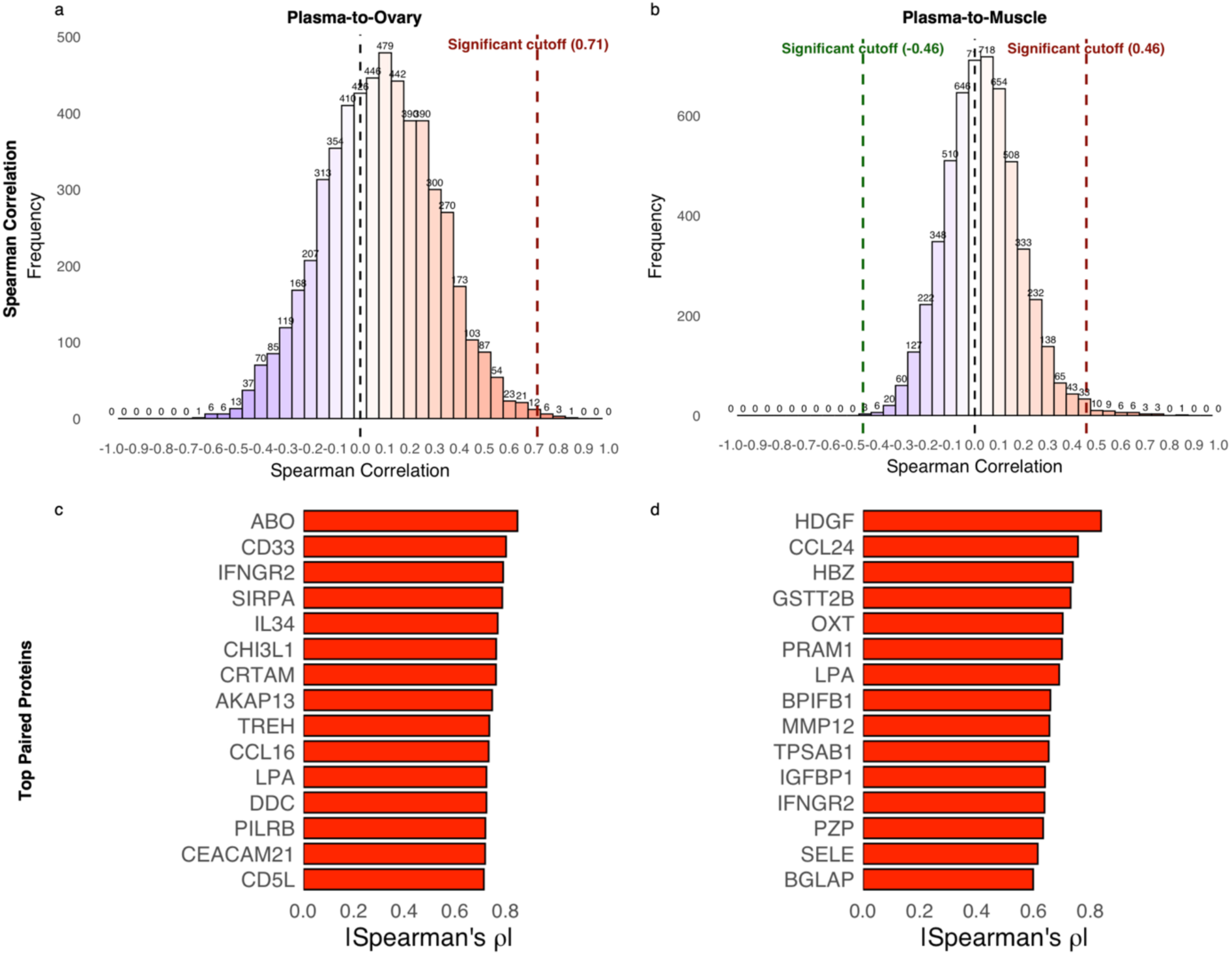
Matched plasma–tissue proteomics identifies a selective subset of tissue proteins reflected in blood. **a**, Distribution of Spearman correlation coefficients for protein abundances between plasma and ovary in 24 matched sample pairs, with significance thresholds indicated. **b,** Same as in a, but for plasma and muscle in 56 matched pairs. **c,** Top plasma–ovary and **d,** top plasma–muscle protein pairs ranked by absolute Spearman correlation above the significance cutoff. Correlations are computed on covariate-adjusted residuals and calibrated against a 1,000-permutation participant-label null; 17 of 5,415 proteins in ovary (median ρ = 0.079) and 50 of 5,415 in muscle (median ρ = 0.028) pass BH-adjusted P < 0.05.

The significantly correlated plasma–tissue protein sets were largely distinct between the two tissues, with 11 proteins observed only for ovary, 44 only for muscle, and six shared between the two tissues (**Supplementary Fig. 3a, b**). The six shared proteins, IFNGR2 (an interferon-receptor component), the immune/metabolic factors IL34 and MBL2, the scavenger receptor CD5L, lipoprotein(a) (LPA) and the carcinoembryonic-antigen family member CEACAM21, were significantly correlated between plasma and tissue in both ovary and muscle, suggesting that they represent conserved tissue-plasma coupling rather than tissue-specific signals. At 80% power with Benjamini-Hochberg control (q = 0.05) across 5,415 assays, the minimum detectable Spearman correlation was |rho| >= 0.77 for the 24 plasma-ovary pairs and |rho| >= 0.54 for the 56 plasma-muscle pairs (**Supplementary Fig. 4**), indicating that only relatively strong plasma-tissue relationships could be detected in the ovarian cohort.

Over-representation analysis (ORA) of Gene Ontology biological processes using the significantly correlated plasma–tissue proteins identified no pathways that remained significant after multiple-testing correction. Similarly, a pairwise analysis testing all cross-protein plasma–muscle associations (each plasma protein against every muscle protein, BH-corrected) recovered only two significant pairs, both negative: plasma GYPA with muscle SERPINF1 (ρ = −0.72) and plasma UCK1 with muscle INPP1 (ρ = −0.74) (**Supplementary Table 3**). Few cross-protein associations were detected under this much larger multiple-testing burden.

### Age-correlated proteomic patterns differ across sex and tissue strata

The number of age-correlated proteins varied markedly across tissues and sexes, with 710 in female plasma, 196 in male plasma, 88 in female muscle, 45 in ovary and only one in male muscle (**Fig. 3a and Supplementary Table 4-8**). Most age-correlated proteins were unique to their respective tissue and sex, with overlap between contexts being limited, suggesting context-dependent age associations, although differences in statistical significance do not themselves establish differences between strata. Female and male plasma differ in sample size, in the age distribution of participants, and in whether batch correction was applied. We therefore explored whether the difference in the number of age-correlated proteins between the two strata is confounded by these factors. To this end, we performed additional down-sampling analysis in the female plasma stratum (**Supplementary Fig. 5a-d and Supplementary Table 9**). At a matched sample size of 23 participants, where both strata were resampled, male plasma yielded a median of 111 age-correlated proteins relative to 13 in female plasma, and a randomly drawn male replicate exceeded a randomly drawn female replicate in 91% of pairs. The age structure of the male stratum contributed in two opposing directions. Sampling from two age extremes increased the number of proteins detected: imposing the male age windows on female plasma at n = 27 under a balanced 13 : 14 allocation returned a median of 90.5 proteins, compared to 29.5 under the age distribution of the female cohort. The 6 : 21 allocation of the male stratum reduced the same comparison to 4, indicating that the imbalance between age groups outweighs any advantage conferred by the extreme age windows (**Supplementary Fig. 5c, d**). Collection site and batch correction accounted for a substantial part of the remaining difference. Female plasma resampled at n = 27 returned a median of 30 proteins across all sites following ComBat correction, 46 across all sites without correction, and 114 when restricted to Wake Forest University, the single site from which all male participants originate. Relative to the male observed count of 196, 6 of 500 female replicates reached this value under the main design (empirical P = 0.014), while 74 of 500 did so in the site-restricted arm (P = 0.150). The direction of the difference was therefore reproducible under every matched design considered, though discovery counts do not themselves measure biological age or aging rate, as estimated by population-scale plasma proteomic models^20^. However, at the present cohort size the difference is only partly separable from collection site and batch correction, and we do not interpret it as a purely sex-specific effect. Direct comparison with mass-spectrometry studies of aging human muscle is complicated by design rather than by biology. Ubaida-Mohien et al. reported 1,265 age-correlated proteins (nominal P < 0.05; 917 at BH-adjusted P < 0.1) among 4,036 proteins quantified in 58 participants aged 20–87 yr, but did so by fitting a single mixed-effects model to both sexes with sex entered as a covariate^17^. Our sex-stratified design halves the effective sample size per test (34 female and 23 male muscle participants), tests 34% more analytes (5,415 versus 4,036), and applies a stricter threshold (BH-adjusted P < 0.05 versus 0.1). Their signal was moreover dominated by mitochondrial, ribosomal and spliceosomal proteins, — intracellular machinery that an affinity-based panel developed for circulating analytes samples only sparsely. The comparison therefore bounds what our muscle arm can detect; it does not indicate an absence of muscle aging in this cohort.

**Fig. 3:**
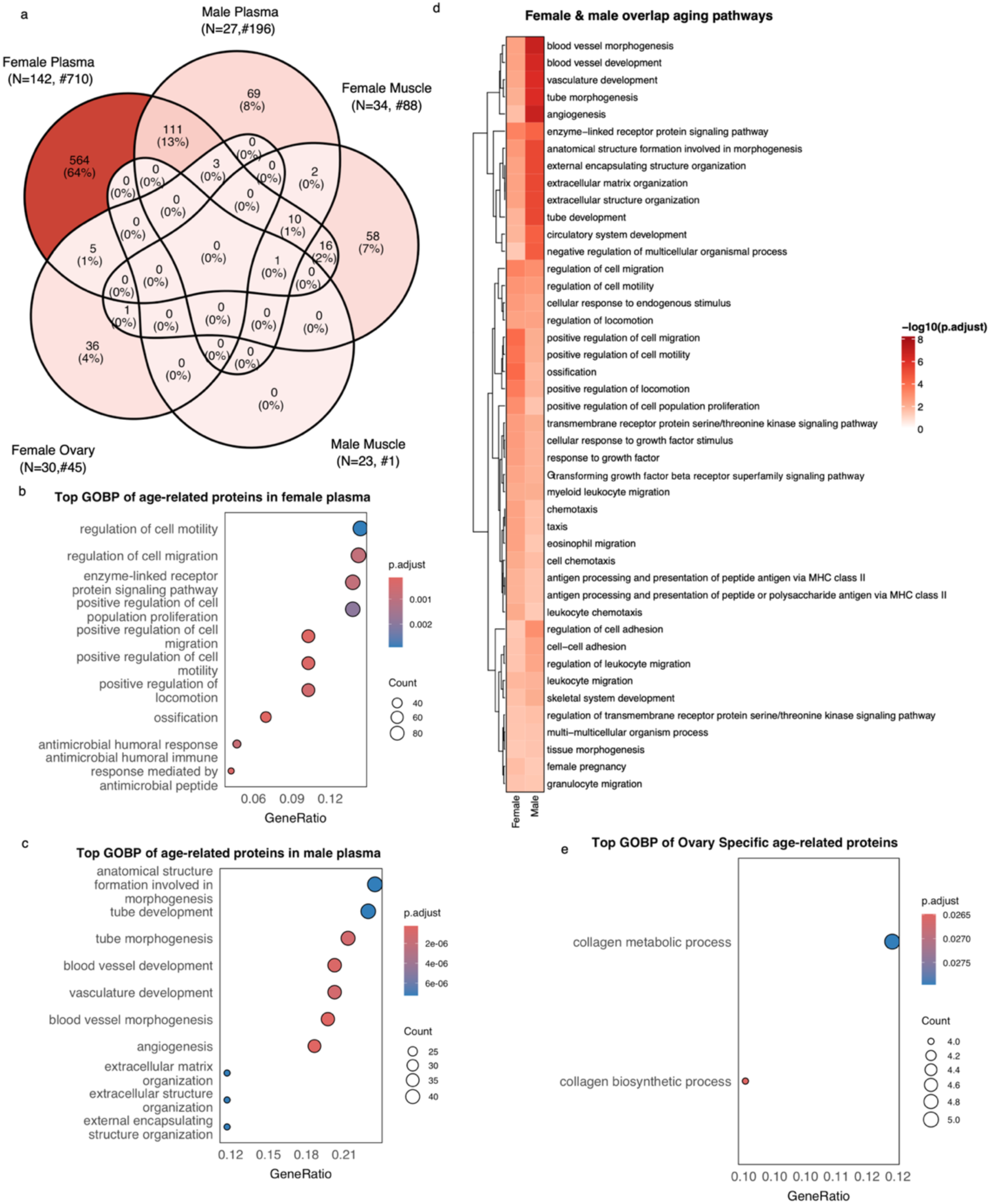
Age-correlated proteomic patterns differ across sex and tissue strata. **a**, Overlap of age-correlated proteins across sexes and tissues. **b,** Female and **c,** male age-correlated proteins in plasma subjected to over-representation analysis (ORA) of Gene Ontology (GO) biological processes; enrichment significance was computed using a hypergeometric test and adjusted for multiple comparisons using the Benjamini–Hochberg method. **d,** Comparison of aging pathways shared between female and male plasma. **e,** GO biological processes enriched among ovary-specific age-correlated proteins, identified by ORA with hypergeometric testing and BH correction.

To characterize sex-stratified aging features within plasma, we performed Gene Ontology ORA separately on the age-correlated proteins identified in female and male plasma. Female-plasma age-correlated proteins were enriched for immune and inflammatory processes, including MHC class II antigen processing and presentation, eosinophil chemotaxis and migration, and antimicrobial humoral response, ossification/bone-growth, and positive regulation of cell migration (**Fig. 3b and Supplementary Table 10**). Chronic low-grade inflammation is the most established systemic feature of human aging, first framed as inflamm-aging^21^ and since linked to cardiovascular disease and frailty^22^. The female plasma signature is compatible with that framework. In contrast, male-plasma age-correlated proteins were enriched for angiogenesis, blood-vessel/vasculature development, extracellular-matrix organization, IGF-receptor signaling and macrophage cytokine production (**Fig. 3c and Supplementary Table 11**). Despite limited overlap in age-correlated proteins, overlap in enriched biological pathways was even more restricted, with only a small set of aging pathways shared between female and male plasma (**Fig. 3d**). Ovary-specific age-correlated proteins were enriched for collagen metabolic and biosynthetic processes (**Fig. 3e and Supplementary Table 12**), whereas the few proteins age-correlated in both muscle and ovary were enriched for developmental and neuro-glial differentiation processes (**Supplementary Fig. 6**). Age-correlated female-muscle proteins, despite numbering 88, yielded no significantly enriched GO term after multiple-testing correction, and male-muscle (n = 1) was not amenable to enrichment analysis. Together, these findings identify different enrichment patterns in the female and male plasma strata, with female plasma characterized primarily by immune and inflammatory processes and male plasma by vascular and extracellular-matrix remodeling, while ovarian aging is dominated by collagen-related pathways, consistent with the progressive fibrotic remodeling of the aging ovary^23–25^.

### LASSO analysis of plasma–tissue observability

Given the generally weak matched-protein correlations, we next explored whether combinations of plasma proteins carried additional information about tissue protein abundance (**Fig. 4a**). Using LASSO regression with plasma proteins as predictors and individual tissue proteins as targets, with covariate residualization (ovary adjusted for age and BMI; muscle adjusted for age, sex and BMI) and imputation performed inside a cross-validation loop so that held-out participants never contribute to their own features or targets, most tissue proteins were not reliably estimated from plasma at these sample sizes. However, a subset showed positive cross-validated associations, comprising 131 ovarian and 164 muscle proteins with cross-validated R² > 0 and BH-adjusted P < 0.05 (**Fig. 4b-c; Supplementary Tables 13–14**). To assess how many of these proteins would be recovered in the absence of a true plasma-tissue relationship, we shuffled tissue participant labels before covariate residualization and performed 100 permutations per tissue using the abbreviated pipeline described in Methods. Both counts exceeded the resulting permutation null (ovary: 131 observed relative to a null mean of 39.2 (s.d. 23.6); muscle: 164 relative to 45.1 (s.d. 21.1); empirical P = 0.0099 in both tissues, the smallest value attainable with 100 permutations), although this comparison used an abbreviated pipeline and no permutation reached either observed count, with a null maximum of 108 in both tissues (**Supplementary Fig. 7; Supplementary Tables 15–16**). The top-ranked predictable proteins were FOLR3 in muscle (R² = 0.84) and BLVRB in ovary (R² = 0.61) (**Fig. 4b, c**). Predictability and tissue of origin were, however, largely decoupled. The best-predicted proteins in muscle were not muscle-enriched at the transcript level: FOLR3 (R² = 0.84), CCL24 (R² = 0.61) and CHIT1 (R² = 0.46) reach their highest GTEx expression in immune tissue, 30- to 800-fold above skeletal muscle, and HDGF (R² = 0.66) is broadly expressed with no organ preference (**Supplementary Tables 17**). Across the 161 predictable muscle proteins with a GTEx match, out-of-sample R² was unrelated to muscle specificity (Spearman ρ = −0.03, P = 0.69), and the set as a whole was not enriched for muscle-origin expression relative to the measured background (median log₂ fold-change muscle-versus-other organs −3.71 versus −4.04; one-sided Mann–Whitney P = 0.087) (**Supplementary Fig. 8a, b and Supplementary Tables 17and 19**). The corresponding ovarian shift was nominally significant (−1.68 versus −2.31; P = 0.025), but both medians remain strongly negative and no ovary-predictable protein was more than two-fold ovary-enriched (**Supplementary Table 18**). In these exploratory models, stronger associations generally involved proteins that were not tissue-enriched at the transcript level. Against that background, two bona fide muscle-restricted proteins — the myosin MYH1 (R² = 0.23) and the metabolic enzyme IDI2 (R² = 0.08) — were nonetheless quantitatively predictable, and were the only proteins in either tissue to exceed the four-fold tissue-enrichment threshold, making them the sole leakage-type candidates this design recovers (**Supplementary Tables 17**).

**Fig. 4:**
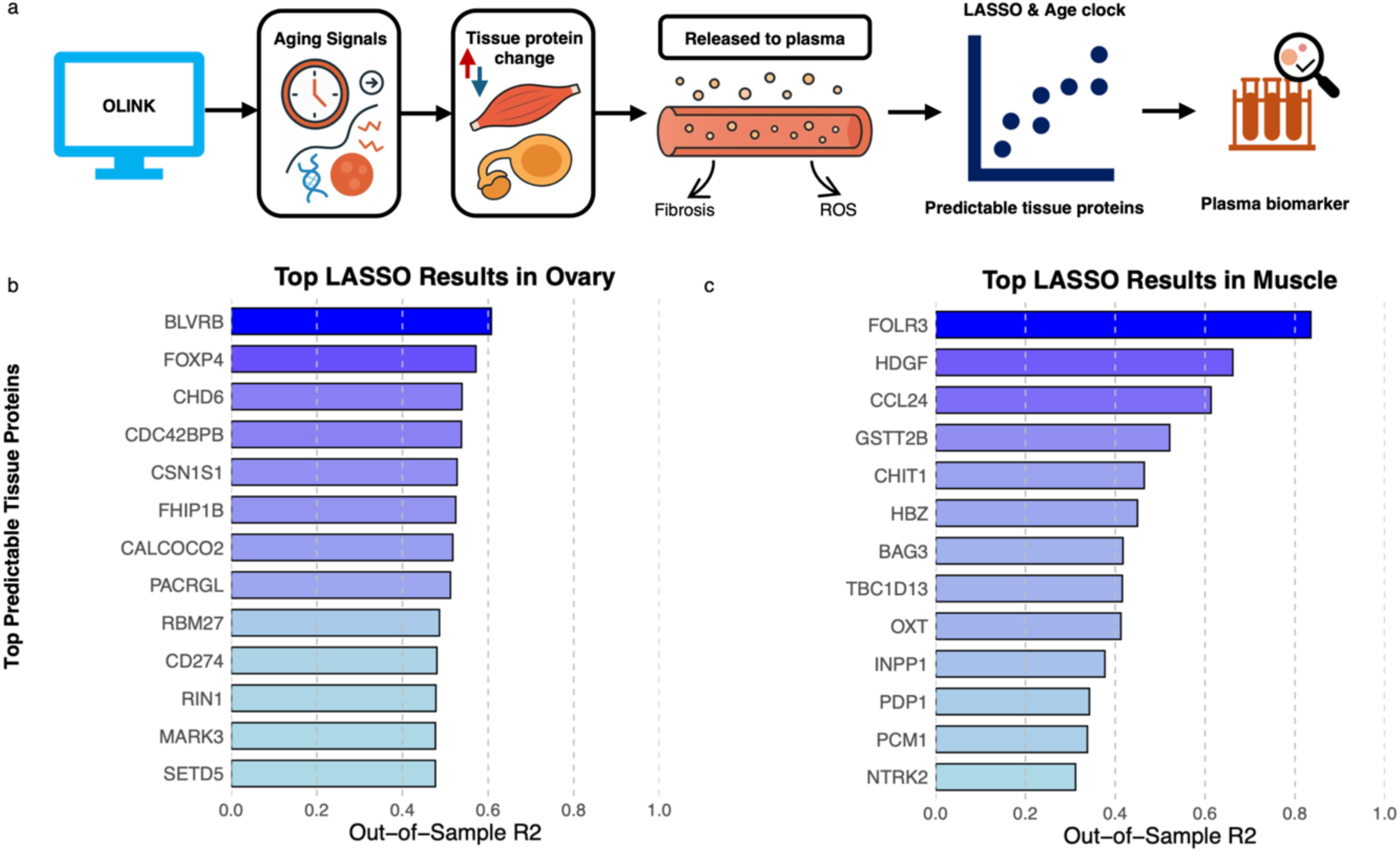
Exploratory multivariate modeling of plasma–tissue relationships. **a**, Schematic of the exploratory plasma–tissue analysis: combinations of plasma proteins are evaluated for information about tissue protein abundance, generating candidates for subsequent biomarker validation. **b–c,** Top-performing tissue proteins ranked by out-of-sample R² for ovary (b; top, biliverdin reductase B / BLVRB, R² = 0.61) and muscle (c; top, folate receptor gamma/FOLR3, R² = 0.84); bar colour encodes R².

We next compared these findings with the Human Protein Atlas single-cell expression atlas (**Supplementary Fig. 9 and Supplementary Table 20**). Among proteins with a single-cell type-of-origin annotation (n=4,840), plasma-legible (concordant, n=57) proteins showed a non-significant tendency toward immune-cell enrichment (odds ratio=1.83, 95% CI 0.82–3.71, uncorrected p=0.083; BH FDR=0.65). No cell lineage was significantly enriched in either the concordant or LASSO-predictable protein sets. The only test whose confidence interval excluded unity was a depletion of neural-origin proteins among the predictable set (OR 0.60, 95% CI 0.37–0.93, P = 0.020, FDR = 0.31), consistent with brain-derived proteins being poorly represented in plasma, but no lineage was significantly enriched or depleted in either set after correction across the 16 tests. The same cell-type-of-origin logic, applied to over 7,000 plasma proteins in 60,542 individuals, supports biological age estimates for more than 40 distinct cell lineages^26^. Our test is a small, underpowered instance of that design, eight collapsed lineages, 57 concordant proteins and 4,840 annotated background proteins, and should be read as exploratory. These exploratory results do not establish that plasma legibility is independent of cell of origin. In a secondary exploratory analysis, a LASSO-based plasma age model showed an association with chronological age (leave-one-out cross-validation; female plasma, n = 142; r = 0.94, R² = 0.87, mean absolute error ≈ 5.15 years; **Supplementary Fig. 8c and Supplementary Table 21**). Because age-preserving ComBat adjustment preceded cross-validation, these performance estimates may be optimistic and do not establish independent age-prediction accuracy. Because the male plasma cohort (n = 27) was too small to train and independently evaluate a separate model, clock performance in males could not be assessed.

We next turned to age-related patterns in female plasma reproductive-axis proteins. Because all ovarian tissue donors were post-menopausal, we examined whether female plasma could detect the menopausal transition by modelling reproductive-axis proteins against age using BMI-adjusted spline regression (FSHB: steepest change at 51 years, bootstrap 95% CI 48–54; age-global FDR = 4×10⁻^33^; **Supplementary Fig. 10, Supplementary Table 22 and Methods**), consistent with the median age at natural menopause reported in US population cohorts (52.54 years in the Study of Women’s Health Across the Nation)^27^. Self-reported menopausal status was not available for these participants, so this correspondence is a cohort-level observation consistent with established endocrine physiology after ComBat adjustment, not per-individual menopausal staging.

Together, these exploratory analyses suggest that, at the sample sizes studied, plasma contains some information to characterize age-related patterns and estimate a subset of predominantly secreted tissue proteins, as well as a few tissue-restricted candidates; larger datasets could reveal additional relationships.

### Aging asymmetrically alters within-pathway protein-rank heterogeneity

Mean shifts in protein abundance capture average aging effects but can miss changes in how proteins act together. We therefore asked whether aging also reorganizes coordination among proteins. Here, coordination denotes concordance of within-pathway protein ranks; EVA does not directly measure regulatory interactions or senescent-cell abundance. Expression Variation Analysis (EVA) quantifies this coordination by comparing the rank-ordered expression of proteins within predefined modules across age groups (**Fig. 5a**), using an entropy-like statistic derived from expected Kendall–tau dissimilarity to test whether a module becomes more heterogeneous with age (age-divergent) or more uniformly coordinated (age-convergent)^28–30^. Participants were stratified into young (20–42 years), middle-aged (43–62 years), and old (63–87 years) groups, using the stratum-specific groupings described above (**Table 1**). Modules were defined as GO biological-process protein sets from Ensembl BioMart, restricted to those with ≥ 50% coverage of their members in our 5,415-protein proteome and ≥ 4 measured proteins, and pruned for redundancy by Jaccard overlap, giving 1,354 modules for analysis. Conceptually, EVA distinguishes pathways that become increasingly heterogeneous with age from those that become progressively more coordinated^31,32^.

**Fig. 5:**
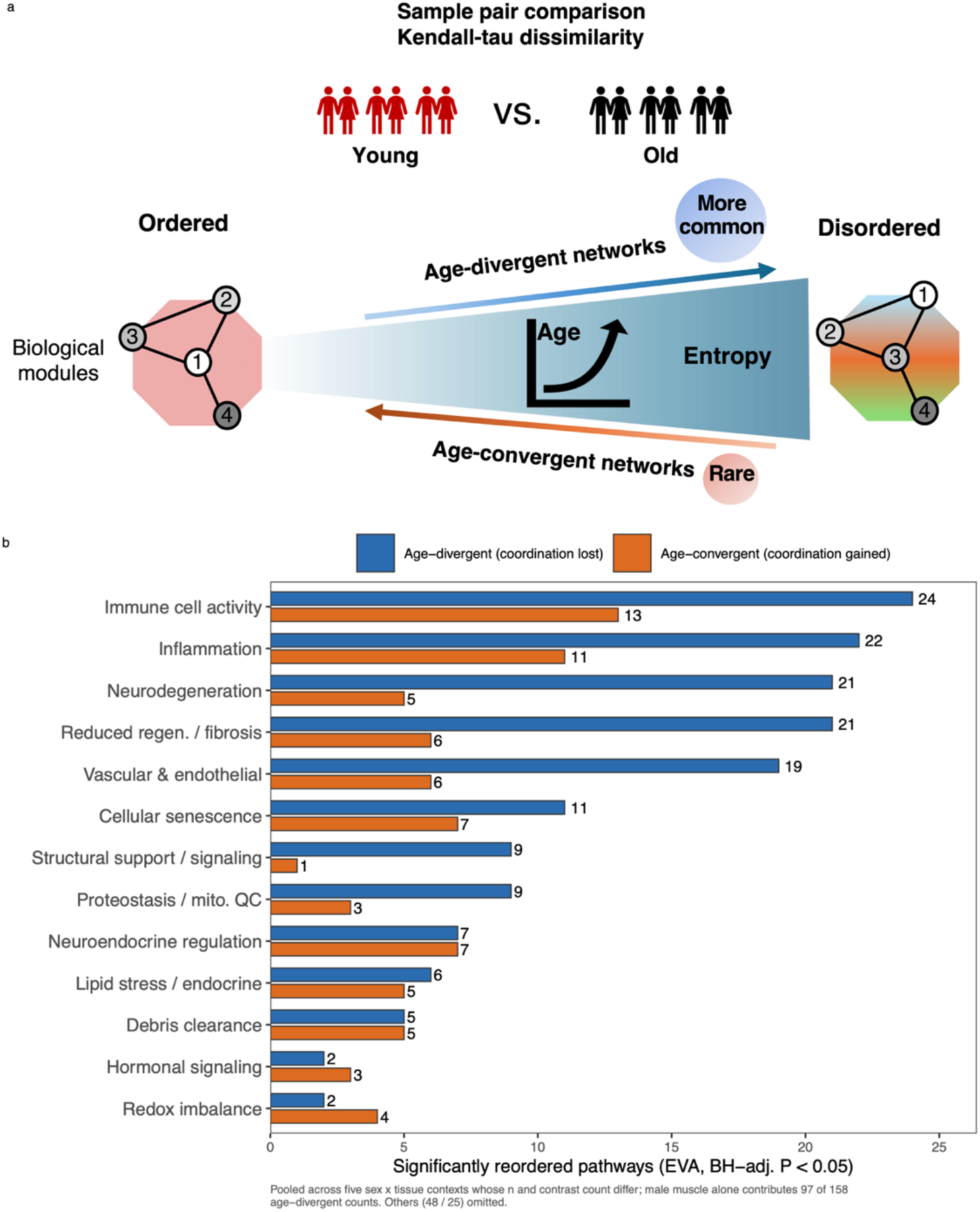
Aging asymmetrically reorganizes protein co-regulation across tissues and sexes. **a**, Schematic of Expression Variation Analysis (EVA). Kendall–tau dissimilarity is computed between pairs of samples within each age group and then compared between groups to quantify differences in within-pathway protein-rank heterogeneity. With age, a module can lose its youthful coordination (age-divergent; blue) or gain coordination (age-convergent; orange). Network diagrams illustrate representative examples of ordered (left) and disordered (right) interaction structures. **b,** Number of significantly reordered pathways per functional module (rows) across the five sex×tissue contexts (columns; F = female, M = male), shown separately for divergent (order lost with age, blue) and convergent (order gained with age, red) reordering; cell values and colour give the count of significant pathways.

**Fig. 5b and Supplementary Table 23** summarizes the number of significantly reordered pathways per module across the five sex × tissue contexts. Network remodeling was asymmetric: 206 pathway-level events were divergent and 101 convergent, indicating that aging in this cohort loosened coordination within modules more often than it tightened it. The full membership of every module-level network is reported by context and direction in **Fig. 6 and Supplementary Tables 24–28** (divergent networks in female plasma, male plasma, female muscle, male muscle and ovary, respectively) and **Supplementary Tables 29–33** (the corresponding convergent networks), while proteins recurring across tissues and sexes are summarized in **Supplementary Table 34 and Fig. 7f**. High-confidence STRING networks (interaction score ≥ 0.7) were resolved for three divergent and six convergent modules in female plasma, four and four in male plasma, four divergent (and no convergent) in female muscle, and a single divergent network in the ovary. Given the broader age structure and larger sample size of the female plasma stratum, we repeated EVA after down-sampling to a matched young-versus-old design (**Supplementary Fig. 11 and Supplementary Table 35-36 and Method**). Convergent reordering remained above chance (18 modules, empirical P = 0.0099), with a similar signal in male plasma (31 modules, P = 0.040), arguing against sample size, contrast number, or ComBat correction as the primary driver.

**Fig. 6:**
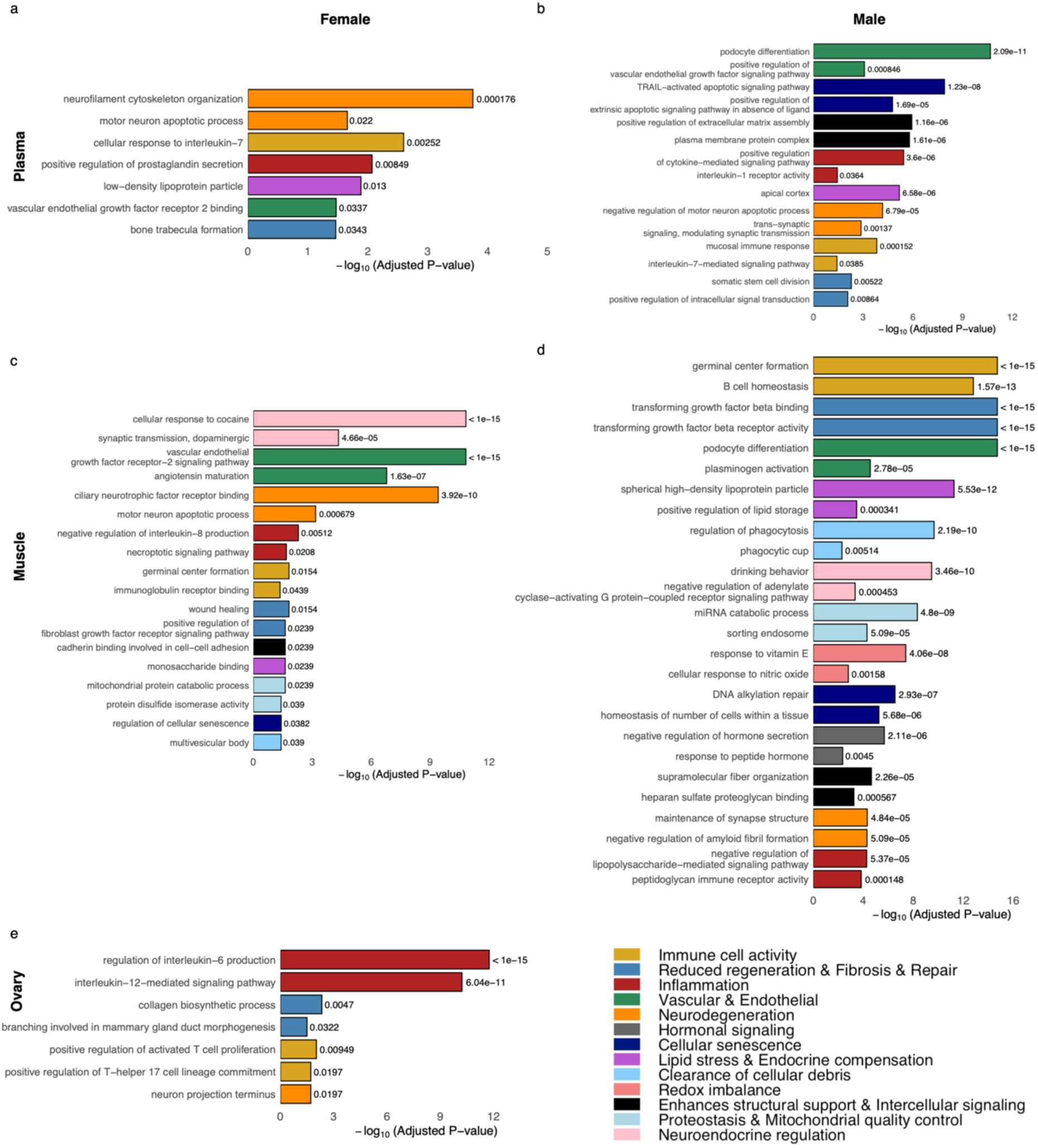
Significantly age-divergent networks across aging in female plasma. **a**, male plasma **b,** female muscle **c,** male muscle **d,** and ovary **e,** as identified by Expression Variation Analysis (EVA). Modules were classified as age-divergent and age-convergent networks based on changes in protein rank coordination, and significance was determined using the Kendall–tau dissimilarity metric with Benjamini–Hochberg correction for multiple testing. These results highlight tissue- and sex-specific rewiring of system-level biological processes during aging.

**Fig. 7:**
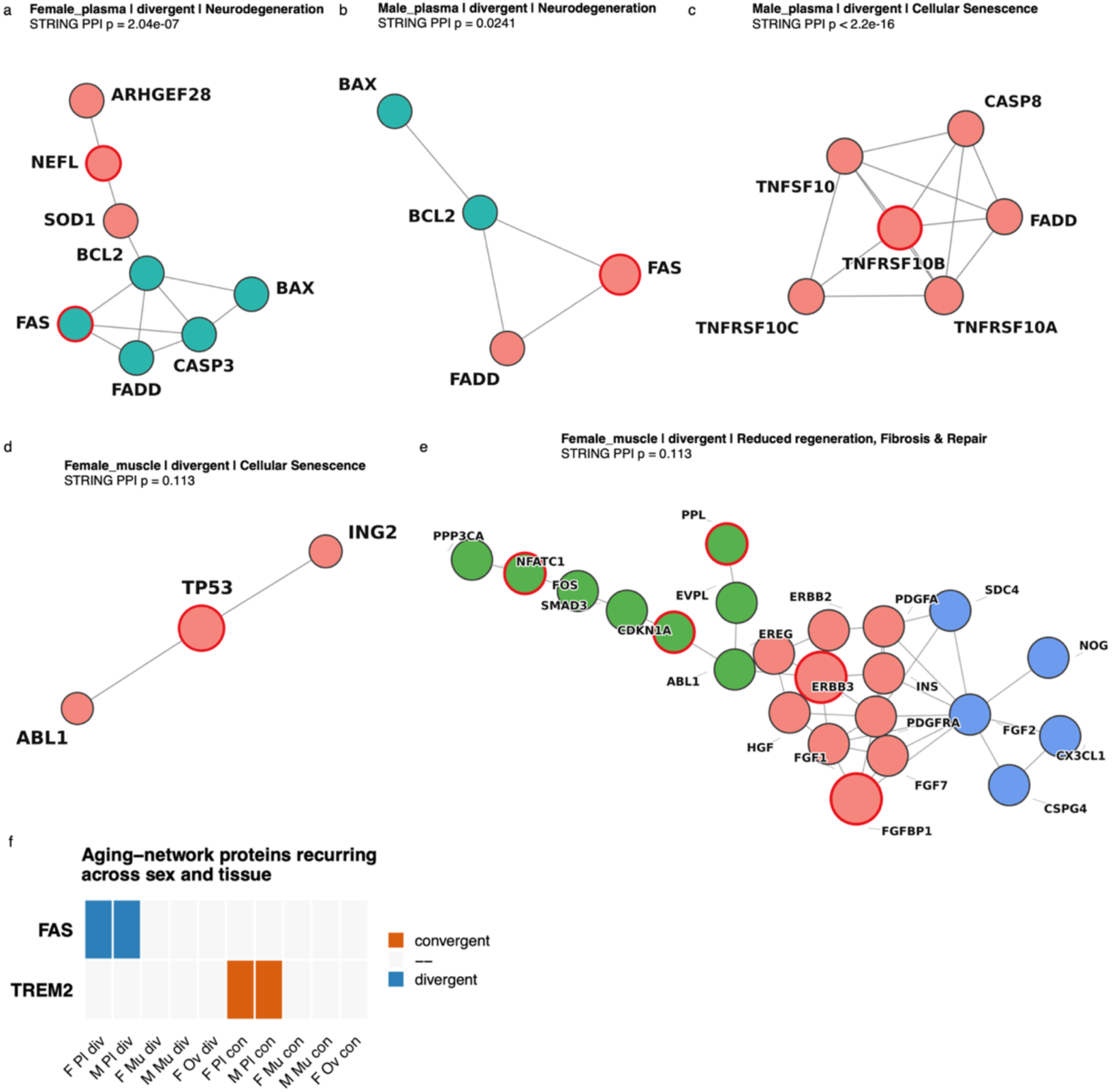
Aging asymmetrically reorganizes interaction networks across tissues and sexes. **a–e**, STRING protein–protein interaction networks of selected age-reordered functional modules. **a,** Age-divergent Neurodegeneration network in female plasma. **b,** Age-divergent Neurodegeneration network in male plasma. **c,** Age-convergent Cellular Senescence network in male plasma. **d,** Age-divergent Cellular Senescence network in female muscle. **e,** Age-divergent Reduced regeneration, Fibrosis & Repair network in female muscle. Node fill denotes STRING MCL sub-clusters, red outlines mark age-associated proteins and edges represent STRING protein–protein associations. **f,** Aging-network proteins recurring across sex and tissue. Columns represent the sex–tissue contexts and direction of reordering in which each protein was identified; coloured cells indicate membership in an age-divergent (blue) or age-convergent (orange) network. FAS recurred in divergent female-plasma and male-plasma networks, whereas TREM2 recurred in convergent female-plasma and male-plasma networks.

Reduced Regeneration, Fibrosis & Repair and Neurodegeneration were the most broadly distributed modules, reordered in nine and eight of the ten context × direction combinations, respectively (**Supplementary Table 23**). The networks resolved from these modules, together with Cellular Senescence, contained overlapping senescence-associated proteins across contexts, consistent with possible roles for senescence-related processes in aging^33–35^. Their membership varied by context, with distinct interaction modules (**Fig. 7a-e**). In plasma, the relevant modules included a death-receptor axis shared by the sexes: the divergent Neurodegeneration networks of female and male plasma were built on the same FAS–FADD–BCL2/BAX module (FAS is the age-correlated protein) (**Fig. 7a, b**), extended in females to CASP3, SOD1 and the neurofilament NEFL (**Fig. 7a**). The same machinery was reordered in the opposite direction in the convergent Cellular Senescence network of male plasma, a complete TRAIL receptor complex (TNFSF10–TNFRSF10A/B/C–CASP8–FADD) and the densest network in the study (6 proteins, 13 edges, density 0.87) (**Fig. 7c**); FADD is the node common to the divergent FAS arm and the convergent TRAIL arm. In female muscle, a related pattern appeared in a divergent TP53–CDKN1A module^17^ (**Fig. 7d**) bridged through ABL1 to the growth-factor core of the largest network we recovered (Reduced Regeneration, Fibrosis & Repair; 23 proteins, 52 edges) (**Fig. 7e**). Together, these findings identify distinct senescence-associated modules in plasma and muscle, with the FAS/FADD/TRAIL death-receptor axis represented in plasma and the TP53–CDKN1A cell-cycle-arrest axis represented in muscle.

The convergent Neurodegeneration networks were organized around different themes in the two sexes: complement- and scavenger-receptor-mediated clearance in female plasma (CLU–C3–ITGAM/ITGB2–LRP1/LRP2–TREM2; **Supplementary Fig. 12a**) and axon-guidance signaling in male plasma (NRP1/NRP2–PLXNA4–SEMA3A/3F; **Supplementary Fig. 12b**). In the post-menopausal ovary, the only significant network was a divergent Reduced Regeneration, Fibrosis & Repair module centered on collagen-associated proteins, consistent with the collagen metabolic and biosynthetic pathways identified among ovarian age-correlated proteins (**Supplementary Fig. 12c**). Of the 67 proteins appearing in any module-level network, only two crossed a sex or tissue boundary: FAS, a member of the divergent networks of both female and male plasma, and TREM2, a member of the convergent networks of both (**Fig. 7f and Supplementary Table 34**). Age-related network remodeling is therefore largely context-specific at the level of protein membership, with death-receptor and myeloid-receptor signaling providing the only cross-sex anchors.

## Discussion

Recent multi-organ proteomic atlases have defined age-associated molecular changes across human tissues and enabled increasingly accurate proteomic aging models ^12,20,36^. Here, we addressed a complementary question: which aspects of tissue aging are represented in the circulating proteome? By profiling matched plasma, skeletal muscle, and ovarian samples from the same individuals using Olink Explore HT, we directly assessed plasma-tissue covariation while minimizing cross-platform technical variation. Muscle and ovary further provided contrasting models of continuous and endocrine aging.

Four principal findings emerged. First, direct plasma-tissue concordance was selective: fewer than 1% of assayed proteins were significantly correlated between plasma and either muscle or ovary. Second, age-correlated proteins and pathways differed substantially across sex and tissue strata. Third, exploratory multivariate models identified additional plasma associations with subsets of tissue proteins beyond matched-protein concordance. Finally, Expression Variation Analysis (EVA) revealed age-associated reorganization of within-pathway protein-rank heterogeneity, including senescence-associated networks in plasma and muscle. Together, these findings position plasma as a selective biologically informative window into tissue aging rather than a comprehensive surrogate for tissue proteomes.

The low direct concordance qualified efforts to infer tissue biology from circulating proteins. Correlations were centered near zero, yet the proteins that concordance included secreted factors and immune receptors such as IFNGR2, IL34, MBL2, CD5L, CEACAM21, CD33, and SIRPA. This pattern is consistent with selective secretion, shedding, or exchange between tissues and circulation, rather than broad representation of intracellular tissue proteins. A related principle has been observed in mice, where plasma metabolomics under lifespan-extending interventions did not preferentially mirror any single tissue^37^. Thus, circulation may be better viewed as a selective sampling compartment whose accessibility depends on protein biology, tissue release and clearance.

Age-associated plasma signatures also differed by sex. Female plasma was characterized primarily by immune and inflammatory processes, whereas the male signature emphasized vascular and extracellular matrix pathways. Population-scale organ-aging models similarly report sex differences that vary across organ systems^13,20^. However, our male cohort was small, and discovery counts were sensitive to cohort structure and collection site. We therefore interpret these results as evidence for differing pathway composition rather than differences in the magnitude or rate of aging between sexes.

Despite limited direct concordance, exploratory LASSO models identified plasma profiles associated with subsets of ovarian and muscle proteins. These associations predominantly involved proteins without strong target-tissue enrichment, consistent with expressed or secreted proteins being more accessible to plasma-based inference. Nevertheless, two muscle-restricted proteins, MYH1 and IDI2, were also predictable and warrant follow-up as candidate tissue-derived biomarkers. Human Protein Atlas analyses provided no significant evidence that plasma-legible or predictable proteins were dominated by a single cellular lineage after multiple-testing correction^26^. Larger matched cohorts will be needed to resolve how tissue provenance, secretion, abundance, and clearance determine plasma legibility.

A secondary plasma age model showed chronological age information is encoded in the circulating proteome, although its performance should be considered exploratory because age-preserving ComBat adjustment preceded cross-validation. Similarly, the midlife FSHB trajectory was consistent with established menopausal endocrine physiology. These analyses support the biological information content of the plasma but were not designed to establish a validated biological-age clock or individual menopausal staging.

EVA provided a complementary perspective by asking how within-pathway protein-rank organization changes with age. Divergent events were more frequent than convergent events (206 versus 101), suggesting that increased within-pathway heterogeneity was more common than increasing coordination in these analyses. Senescence-associated proteins appeared across Neurodegeneration, Cellular Senescence, and Reduced Regeneration, Fibrosis & Repair modules^33–35^. In plasma, FAS-FADD-BCL2/BAX networks diverged with age in both sexes, whereas a TRAIL receptor complex containing TNFSF10, TNFRSF10A/B/C, CASP8, and FADD converged in male plasma. Female muscle instead showed a divergent TP53-CDKN1A module. These findings suggest that related senescence-associated processes can undergo distinct forms of network reorganization across biological contexts, without implying corresponding changed in pathway activity or senescent-cell abundance.

Cross-context recurrence at the protein level was limited: FAS and TREM2 were the only proteins recurring across sex or tissue boundaries in the resolved networks. Thus, age-associated network remodeling largely context-specific in its molecular membership even when broader functional themes recur. This distinction cautions against interpreting a circulating senescence-associated signature as a direct molecular readout of the corresponding tissue state and instead supports network organization as a complementary dimension of proteomic aging.

The ovarian findings provide an additional example of this selectivity. Our post-menopausal ovarian cohort showed a collagen-dominated age-associated program consistent with the stromal fibrosis reported in human ovarian aging atlases^19,38^. Yet this tissue remodeling was largely not directly reflected in matched plasma, reinforcing the distinction between molecular changes occurring within an organ and those accessible through circulation.

Several limitations should be considered. Cohort size was modest, particularly for male and ovarian samples, limiting power to detect weaker and sex-specific associations. The ovarian cohort represents a late-life endocrine window rather than the full reproductive lifespan, and only two tissues were examined. The affinity-based panel also samples a different portion of the proteome than untargeted mass spectrometry, particularly for intracellular proteins. Finally, the cross-sectional design precludes inference about longitudinal trajectories or causality, and the plasma-tissue relationships were not tested in an independent matched cohort. Larger longitudinal studies spanning additional organs, populations, and complementary proteomic platforms will be important for testing generalizability and improving tissue-state inference.

Collectively, our findings support plasma proteomics as a selective molecular window into human tissue aging. Although only a small fraction of assayed tissue proteins showed direct plasma concordance, circulating proteins retained biologically structured information about age-associated pathways, network and multivariate analyses therefore provide a framework for distinguishing aging signals accessible in circulation from molecular changes that remain tissue-specific, and for developing more interpretable circulating biomarkers of human aging.

## Methods

### Participant recruitment and sample collection

Human skeletal muscle, ovarian tissue, and peripheral blood samples were collected under Institutional Review Board (IRB)-approved protocols with written consent from all participants. Skeletal muscle biopsies and matched plasma samples were obtained from healthy adult male and female participants (24-87 years) enrolled through the Cellular Senescence Network Tissue Mapping Center at Wake Forest University under a single IRB-approved protocol (WF IRB00077384). Histologically normal ovarian tissue and matched plasma samples were obtained through the Northwestern University Reproductive Tissue Library under IRB-approved protocols (STU00215770 and STU00215938) from women undergoing gynecological surgery. Additional plasma samples from healthy female participants were obtained through the Susan G. Komen Tissue Bank at the Indiana University Simon Comprehensive Cancer Center. Breast tissue from these participants was not included in the present study^39^.

Peripheral blood was collected at the time of tissue sampling and processed according to standardized protocols. Skeletal muscle biopsies were obtained using a standardized percutaneous vastus lateralis biopsy procedure (PMID: 38315316), while ovarian tissue specimens were collected immediately following surgical excision. Tissue samples were processed immediately after collection, snap-frozen in liquid nitrogen, shipped on dry ice, and stored at −80°C until proteomic analysis. The study included 57 skeletal muscle samples, 30 ovarian tissue samples, and 169 plasma samples, comprising 56 matched plasma-muscle pairs and 24 matched plasma-ovary pairs (**Table 1**).

### Sample processing and proteomic analysis

Frozen skeletal muscle and ovarian tissue samples (20-40 mg) were homogenized in a RIPA lysis buffer supplemented with protease and phosphatase inhibitors using a bead-based tissue homogenizer. Tissue lysates were clarified by centrifugation, protein content quantified by BCA, and normalized to 1 mg/mL. For plasma samples, 50 µL aliquots were prepared without additional processing. Tissue lysates and plasma aliquots were analyzed by Olink Proteomics (Boston, MA, USA), enabling proteomic profiling of both plasma and tissue using the same antibody-based assay^40^.

### Data Preprocessing

All bioinformatics analyses were conducted in R (v4.4.2). Prior to downstream analyses, stringent sample-level quality control (QC) was performed. Samples failing either assay-level or sample-level QC were excluded. Additional outlier detection was carried out using principal component analysis (PCA) and distributional outlier detection (median/IQR) via olink_pca_plot and olink_qc_plot, and samples flagged by both methods were removed. Control samples were also excluded.

After sample QC, proteins with more than 10% missing data across retained samples were removed. For the remaining dataset, missing values were imputed using the Random Forest–based imputation method implemented in the missForest R package (v1.5)^41^.

Subsequent preprocessing diverged depending on the analysis type:

- For all analyses except Expression Variation Analysis (EVA), PC-normalized NPX values were used. Protein expression matrices were further standardized using Z-score normalization via the base R scale function.
- For EVA, raw protein count data were used instead of NPX values. After applying the same QC procedures, no further normalization was performed to preserve intrinsic expression variability.

### Spearman Correlation Analysis

To quantify plasma–tissue correlation, we computed Spearman rank correlations between plasma and tissue abundances for each protein detected in both compartments, using quality-controlled matched pairs (56 plasma–muscle and 24 plasma–ovary). Prior to correlation, each protein was residualized against available covariates by linear regression (age, sex and BMI; covariates with no variation in a given comparison were dropped automatically, so the single-sex ovary comparison was adjusted for age and BMI only). Significance was assessed using a participant-label permutation null rather than the analytic Spearman distribution. Within each tissue, the tissue residual matrix was permuted across participants 1,000 times, and each protein’s observed |ρ| was compared with the pooled null distribution comprising all proteins × permutations. The resulting pooled permutation P values were corrected across all tested proteins within each tissue using the Benjamini–Hochberg (BH) method^42^. Minimum detectable correlation for the matched-pair analyses were estimated using pwr.r.test from the pwr R package^43^, using as the two-sided α the largest raw permutation P value declared significant at BH-adjusted P < 0.05 in each tissue (1.5 × 10⁻⁴ for ovary and 4.5 × 10⁻⁴ for muscle).

### Detectability relative to the limit of detection (LOD)

To assess whether limited assay sensitivity contributed to the low plasma–tissue concordance, we evaluated protein detectability relative to the limit of detection (LOD). Assay-specific LODs were obtained using ‘olink_lod()’ from the OlinkAnalyze R package with the FixedLOD method and the official Olink Explore HT fixed-LOD file. Fixed LODs were used because each delivery contained fewer than the ten negative-control wells required for negative-control-based LOD estimation (four for plasma and two for each tissue run, all from the same control lot). To ensure comparability across compartments, plate-control-normalised LODs were used throughout, consistent with the plate-control-normalised NPX values analysed across all three deliveries. For each protein and compartment, we calculated the detection rate as the proportion of samples with NPX above the LOD and the mean signal above LOD as the mean NPX − LOD. Detectability was evaluated using the same matched participants included in each plasma–tissue correlation analysis: plasma measurements from the 24 ovary-matched participants and 56 muscle-matched participants were evaluated separately, and tissue measurements were evaluated in the corresponding matched participants. The association between detectability and plasma–tissue concordance was quantified by Spearman correlation between |ρ| and mean signal above LOD.

### LASSO regression and plasma proteomic age clock

To systematically identify plasma predictors of tissue protein levels, we used LASSO regression with repeated k-fold cross-validation, implemented with the glmnet package (v4.1-10) in R^44^. For each protein, plasma and tissue abundances were residualized against covariates by linear regression within each cross-validation training partition — age, sex and BMI for the mixed-sex muscle comparison, and age and BMI for the single-sex ovary comparison — so that predictions reflect plasma–tissue coupling beyond shared demographic structure.

The residualized data were subjected to repeated k-fold cross-validation (k = 5 folds, 6 repeats). Covariate residualization and residual missing-value imputation were performed inside the cross-validation loop rather than once on the complete matrix: within every outer training partition the covariate coefficients were fitted on the training participants only and then applied to the held-out participants, so that a held-out sample never contributes to the model used to construct its own predictors or its own target. Missing values occurred only in the plasma matrix (<1% of values); the tissue matrices had none. Within each training partition, the originally missing plasma values were reset to missing and re-imputed with training-partition protein medians, so the missForest values computed on the full matrix were not used by the LASSO. The only steps applied to the full dataset before cross-validation were the protein missingness filter (which removed no proteins) and this superseded missForest imputation. No batch correction or feature pre-selection was performed: all 5,415 plasma proteins entered every model, and predictor selection occurred only through the L1 penalty within training partitions. Within each fold, the optimal L1 penalty (lambda.min) was chosen by internal 5-fold cross-validation (cv.glmnet, alpha = 1) on the training partition, a LASSO model was refit at that penalty, and predictions were generated for the held-out samples; out-of-fold predictions were averaged across the 6 repeats to give one cross-validated prediction per sample. Per protein, predictive performance was summarized as the out-of-sample R² and the Spearman correlation between observed and cross-validated predicted abundance, with significance BH-corrected across proteins^42^. A tissue protein was called predictable from plasma if it had BH-adjusted P < 0.05 and R² > 0. A fixed random seed (seed = 123) was set for reproducibility. For the plasma proteomic age clock, we trained a single LASSO model to predict chronological age from all plasma proteins in a leave-one-out cross-validation loop (cv.glmnet, alpha = 1, 10 inner folds; residual missing values imputed by column median), yielding one leave-one-out age estimate per participant. The clock used the female-plasma matrix after cohort-wide, age-preserving ComBat correction (n = 142); its cross-validation results are therefore descriptive rather than independent out-of-sample validation.

### Permutation null for the plasma-to-tissue LASSO

To establish that the predictable-protein counts exceed what the pipeline returns on data with no plasma–tissue coupling, an abbreviated version of the lasso procedure was evaluated under a participant-label permutation null. In each permutation the tissue participant labels were shuffled before covariate residualization: this breaks the plasma–tissue participant pairing and the tissue–covariate correspondence together. The plasma folds were left untouched and reused, since only the tissue limb of the model is affected by the shuffle. One hundred permutations were run per tissue, each using a single 5-fold outer cross-validation over a fixed panel of 500 targets drawn once and stratified across abundance deciles; counts were then scaled linearly to the full 5,415-protein panel. This abbreviated design was adopted for computational feasibility. Repeated outer cross-validation in the observed run serves to stabilize each protein’s out-of-sample estimate against the choice of fold split, whereas the null only needs to estimate how often a protein passes the predictability criterion by chance, which the 100 permutations already average over. The single outer split was the first repeat of the observed run, so observed and null shared the same fold assignment, and covariate residualization, imputation and lambda selection remained inside the training folds. Because the BH threshold depends on the full p-value distribution, each permutation counted panel proteins with a raw P at or below the P-value cutoff realized by BH in the observed run and R² > 0. As a sensitivity check, the BH correction was also re-run within the 500-protein panel instead of scaling a fixed threshold (scaled null mean 18.2 and 17.2, maximum 108.3 and 75.8, for ovary and muscle, respectively). Because observed predictions were averaged over six repeats whereas each permutation used a single split, the two procedures are not strictly identical; however, no permutation under either counting rule reached the observed counts.

### Age-correlated proteins

Age-correlated proteins were identified within each sex- and tissue-defined context (female plasma, male plasma, female muscle, male muscle and ovary) by Spearman rank correlation between each protein’s abundance and participant chronological age. For every protein, the correlation was computed over all samples with non-missing values in that context (two-sided Spearman test); the resulting p-values were corrected for multiple testing across all tested proteins in the context by the Benjamini–Hochberg method, and a protein was called age-correlated when its BH-adjusted P < 0.05. The correlation direction (positive or negative with age) and the number of samples used were retained for each protein (**Supplementary Tables 4–8**). Age-correlated protein analysis in female plasma was performed on ComBat-adjusted data to account for site-related batch effects.

### Down-sampling sensitivity analysis for age-correlated analysis

The five strata differ in size (female plasma 142, female muscle 34, ovary 30, male plasma 27 and male muscle 23 participants) and, in the male arms, in how participants are distributed across age, so the raw discovery counts are not directly comparable. Two resampling analyses were therefore performed. First, every stratum was down-sampled at random, without any age balancing, to a common N = 23 (the size of the smallest stratum) over 500 replicates, and to a shared grid of sample sizes over 200 replicates per grid point. Within each replicate, age-correlated proteins were recounted exactly as in the full analysis (Spearman correlation with age, BH-adjusted P < 0.05).

Second, because the male participants were sampled at two age extremes rather than continuously, female plasma was additionally down-sampled to the male plasma stratum’s own design. The male age windows were read from the male participants themselves and are 23–30 and 70–87 years for plasma (6 young : 21 old, n = 27) and 23–30 and 70–82 years for muscle (5 young : 18 old, n = 22). Female participants were drawn from inside those windows at the male allocation, and an allocation-balanced variant (13 : 14) was run alongside to separate the effect of the extreme-age design from the effect of the allocation. A site-restricted arm, using only female plasma participants from Wake Forest University without ComBat correction, was included so that the comparison could also be made within the single site from which all male participants originate. The same design was applied to female muscle against the male muscle windows.

### Over-Representation Analysis (ORA)

ORA was used to test whether Gene Ontology biological processes (BPs) are enriched relative to the study background. In this study, we applied ORA separately to (i) significantly correlated protein pairs from the plasma–ovary and plasma–muscle analysis, (ii) sex-specific and (iii) tissue-specific age-correlated protein sets.

The significant proteins were first mapped from UniProt identifiers to gene symbols using the Bioconductor package ‘clusterProfiler’ (v4.14.6) with the annotation database ‘org.Hs.eg.db’ (v3.20.0). Subsequently, Gene Ontology (GO) biological process enrichment was performed using the ‘enrichGÒ function from the ‘clusterProfiler’ package^45^. The background universe for enrichment analyses comprised all gene symbols detected in the complete proteomic dataset. Statistical significance for enrichment was determined using the hypergeometric test (equivalent to a one-sided Fisher’s exact test), and multiple-testing correction was applied using the BH method^42^. BPs with BH-adjusted p-values < 0.05 were considered significantly enriched.

### Batch correction of female plasma samples

To correct for site-specific batch effects present only in the female-plasma data, we applied ComBat from the sva package (v3.54.0) in R^46,47^. Sampling site was specified as the batch variable, and age and BMI (both z-scored) were retained as biological covariates in the model matrix to preserve age- and BMI-related structure. The ComBat-corrected expression matrix was used for all downstream analyses involving female plasma, including age-correlated protein discovery and EVA network analysis. The matched plasma–tissue analyses (Spearman correlation and LASSO) used plasma data without ComBat correction: all matched ovary participants were sampled at Northwestern University and all matched muscle participants at Wake Forest University, so site was constant within each analysis and no batch correction was required.

Batch correction markedly reduced the proportion of proteins associated with site-specific variation — from 38.449% in the age-analysis input matrix and 37.285% in the EVA input matrix before correction to a negligible fraction afterward — without diminishing the variance attributable to age and BMI, as verified by variancePartition before and after correction^48^ (**Supplementary Fig. 1**).

### Expression Variation Analysis (EVA) Network Analysis

EVA was performed to identify biological pathways exhibiting altered coordination structures across age groups. Unlike conventional methods that evaluate mean expression differences, EVA focuses on within-group variability by computing Kendall–tau dissimilarity across all sample pairs. For a given pathway, EVA defines an entropy-like statistic *E* as the expected rank discordance between two samples:

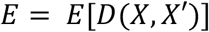

Here, *X* and *X*^!^ are two independently sampled individuals from the same phenotype group, and *D*[*X*, *X*^!^] denotes the Kendall–tau distance, which quantifies the number of pairwise order violations in protein expression rankings within the pathway. Higher *E* values indicate greater variability in gene–gene coordination, reflecting a more disordered network structure.

Protein data input for EVA consisted of raw, unstandardized counts obtained from Olink Explore HT (NPX Explore v7.2.1) proteomics measurements. EVA was performed with the Bioconductor package GSReg (v1.40.0). Functional modules were defined as Gene Ontology biological-process protein sets obtained from Ensembl BioMart (v2.62.1)^49^ and mapped onto the 5,415 measured proteins; modules were retained only if they covered ≥ 50% of their annotated members in our proteome and contained ≥ 4 measured proteins, and near-duplicate modules were pruned by Jaccard overlap, leaving 1,354 modules tested. For each module, EVA compared age-group contrasts appropriate to the stratum (female plasma: Young vs. Middle-age, Young vs. Old, and Middle-age vs. Old; male plasma, female muscle and male muscle: young vs. old; ovary: middle-age vs. old). For female plasma, site-corrected (ComBat-adjusted) log scale matrices were used; other strata used raw counts from their single site without further normalization. Statistical significance was assessed by permutation testing with Benjamini–Hochberg correction^42^, and modules with BH-adjusted P < 0.05 were considered significantly reordered.

We further classified these pathways into two descriptive categories:

- Age-divergent networks: modules with significantly increased entropy in the older group (i.e., E_younger_ < E_older_), indicating greater within-pathway protein-rank heterogeneity.
- Age-convergent networks: modules with decreased entropy in the older group (i.e., E_younger_ > E_older_), reflecting lower within-pathway protein-rank heterogeneity.

These two categories represent distinct patterns of network remodeling across age and were used to quantify the directional impact of aging on proteomic regulation.

### Permutation null and design-matched down-sampling for EVA

To assess whether results in female plasma reflected differences in sample size, age-group composition or site-related effects, we performed two down-sampling analyses. The first assessed sample-size and age-design effects by matching female plasma to the male plasma design, whereas the second additionally assessed site and ComBat effects by restricting the analysis to the collection site shared by both sexes and using uncorrected data. In the first down-sampling analysis, 6 young and 21 old female participants were sampled from within the corresponding male age ranges (23–30 and 70–87 years, respectively; n = 27) in each of 100 replicates, and EVA was repeated using a single young-versus-old contrast. In the second down-sampling analysis, the same sampling scheme was applied to uncorrected female plasma from Wake Forest University, with 100 replicates. The number and identity of modules were recorded for each replicate.

Because sex-specificity is defined elsewhere in this study as a set difference rather than as a count, the down-sampled EVA was also evaluated at the level of module identity: for every replicate we recorded the overlap with the male convergent set, the Jaccard index, the female-only and male-only counts, the overlap expected by chance for sets of those sizes, and two-sided hypergeometric P values for enrichment and for depletion of the overlap. A module was called recurrent when it was significant in at least 50% of the replicates that tested it, the denominator being the number of replicates in which GSReg actually returned that module rather than the total number of replicates.

### STRING Protein-Protein Interaction (PPI) Network Analysis

To elucidate functional relationships and identify key protein interactions within age-correlated modules identified by EVA, we performed PPI network analyses using the STRING database (v12.0)^50^. Samples were first stratified by sex and tissue, and EVA was then applied within each stratum to detect age-convergent and age-divergent networks. These networks were subsequently categorized into one of 12 biologically meaningful functional groups spanning Immune cell activity, Reduced regeneration & fibrosis & repair, Inflammation, Neurodegeneration, Hormonal Signaling, Cellular Senescence, Lipid Stress & Endocrine Compensation, Clearance of Cellular Debris, Redox Imbalance, Enhances Structural Support & Intercellular Signaling, Proteostasis & Mitochondrial Quality Control, Neuroendocrine regulation. When the number of networks within a group was insufficient for meaningful categorization, all networks were analyzed collectively without further subdivision. STRING analyses were independently conducted for each functional group or combined network set, using high-confidence co-expression-based interactions (interaction score > 0.7) linking at least one age-altered protein to others. Protein clustering was performed using the Markov Cluster Algorithm (MCL)^51^, and disconnected nodes (degree = 0) were excluded from the final visualizations.

### Plasma Biomarker Selection and GTEx analysis

To identify candidate proteins suitable for plasma biomarker, we integrated the identified age-correlated proteins with predictive modeling results from cross-validated LASSO regression analyses. Proteins in tissue that were significantly age-correlated and showed exploratory multivariate associations with plasma profiles were treated as candidates for follow-up. These proteins are exploratory biomarker candidates requiring validation before they can be interpreted as non-invasive monitors of tissue-specific aging changes.

To determine whether a predictable plasma signal reflects tissue-private biology or spillover from more highly expressing organs, we performed a GTEx tissue-of-origin analysis for every predictable protein. Median transcripts-per-million (TPM) values from GTEx v10^52^ were collapsed from sub-tissues to organs by taking the maximum TPM per organ, and each protein was scored for tissue enrichment as the ratio of expression in the target organ to the maximum across all other organs; a protein was called tissue-enriched when this ratio was ≥ 4-fold and target expression was ≥ 1 TPM. This four-fold criterion follows the organ-enrichment definition used to construct plasma-based organ aging models^13^, so that our tissue-of-origin assignments are directly comparable to that literature. The log₂ fold-change of target-vs-other-organ expression was compared between predictable proteins and the measured background by a one-sided Mann–Whitney U test. Predictable, age-correlated tissue proteins that were also tissue-enriched in GTEx were prioritized as candidate ‘leakage’ biomarkers.

### Cell-type-of-origin analysis (Human Protein Atlas)

To determine the cell type of origin of the plasma-legible proteins, we annotated every panel protein with its most cell-type-specific single-cell expression from the Human Protein Atlas (proteinatlas.tsv; the ‘RNA single cell type specific nCPM’ field)^53,54^, taking the top-ranked cell type per gene symbol and collapsing it to one of eight cell lineages (immune, endothelial, epithelial, muscle, fibroblast/stromal, neural, germ/reproductive, or other secretory) by keyword mapping; unannotated proteins were excluded so as not to dilute the tests. The background was the set of panel proteins carrying such an annotation (n = 4,840). Against this background we tested two gene sets — the plasma–tissue concordant set (proteins significant in the muscle or ovary pairwise analysis, n = 57) and the broader LASSO-predictable set (BH-adjusted P < 0.05 and R² > 0 in muscle or ovary, n = 251) — for enrichment or depletion of each lineage using a two-sided Fisher’s exact test per lineage and set, reporting the odds ratio with 95% confidence interval, the set and background counts (k, n), and a Benjamini–Hochberg FDR computed across all 16 tests. In parallel, as a transcription-factor landscape layer, each protein was flagged as a transcription factor from the HPA ‘Protein class’ field; we quantified the fraction of the annotated panel that are transcription factors (4.3%; 207/4,840).

### Reproductive-axis protein trajectories

To test whether the menopausal endocrine transition is legible in the circulating proteome, eight reproductive-axis candidate proteins (FSHB, LHB, SHBG, PRL, INHBB, INHBC, WFDC2 and MUC16) were modeled as a function of age in female plasma only, on the ComBat-adjusted, standardized expression matrix. For each protein, abundance was regressed on age with a natural cubic spline (ns, 3 degrees of freedom) and BMI as a covariate; the fitted trajectory was evaluated at the median BMI and displayed with its pointwise 95% confidence band (fit ± 1.96 × SE). The overall effect of age was assessed by comparing the spline model with a BMI-only null model using a nested-model F-test, and departure from linearity by comparing the spline model with a linear-age model; p-values were Benjamini–Hochberg-corrected across the eight candidates. As a complementary group-level analysis, samples were binned into young (≤42 y), middle-aged (43–62 y) and old (>62 y) groups and compared by Kruskal–Wallis test followed by Dunn’s post-hoc test (Benjamini–Hochberg-adjusted). The age of steepest change was estimated as the location of the maximal absolute first derivative of the fitted spline, restricted to the interior (5th–95th percentile) of the observed age range, with a bootstrap 95% confidence interval (2,000 resamples). This change point was considered identifiable only when its bootstrap interval was sufficiently narrow (width ≤ 40% of the age span); among the eight candidates the criterion was met only for FSHB, which is shown in Supplementary Figure 10 (steepest change at 51 years, bootstrap 95% CI 48–54).

## Data Availability

The Olink Explore proteomic data (normalized protein expression (NPX) values) generated in this study from plasma, muscle and ovary samples, together with the processed protein-abundance matrices and sample metadata, have been deposited in Zenodo (https://doi.org/10.5281/zenodo.23428018). Source data for all main and supplementary figures are provided in Supplementary Tables 1–36.

## Code Availability

All custom code used for data processing, statistical analyses and figure generation is available on GitHub (https://github.com/qiongronghuang/plasma-tissue-protein-aging) and has been archived in Zenodo (https://doi.org/10.5281/zenodo.23428018).

## Acknowledgements

We gratefully acknowledge the women and men who generously donated ovarian tissue, skeletal muscle tissue, and plasma for this study. Their contributions made this work possible. Plasma samples from the Susan G. Komen Tissue Bank at the IU Simon Comprehensive Cancer Center were used in this study. We thank the contributors who collected and provided samples used in this study, including Indiana University, as well as the donors and their families for their invaluable participation. This work was supported by the National Institutes of Health (NIH) Common Fund Cellular Senescence Network (SenNet; Office of the Director) program through NIA grant U54 AG075932 (PI: Schilling; MPI: Melov).

## Contributions

Concept and design: Birgit Schilling, Nathan D. Price, Qiongrong Huang, John C. Earls, Noa Rappaport, Mark A. Watson. Analysis or interpretation of data: Qiongrong Huang, John C. Earls, Noa Rappaport. Drafting of the manuscript: Qiongrong Huang, Mark A. Watson. Data collection: Mark A. Watson, Heather L. Gregory, John Stone, Maryellen G. Pavone, Jamie N. Justice, Bumsoo Ahn, Ryan P. Lamers, Christopher C. Benz. Combat code: Qiongrong Huang, Annalise Schweickart. Critical revision of the manuscript for important intellectual content: All authors. Funding acquisition: Birgit Schilling, Francesca E. Duncan.

## Supervision

John C. Earls, Birgit Schilling, Noa Rappaport, Francesca Duncan, Nathan D. Price. All authors approved the final version of the manuscript for submission.

---

**Supplementary Figure 1:**
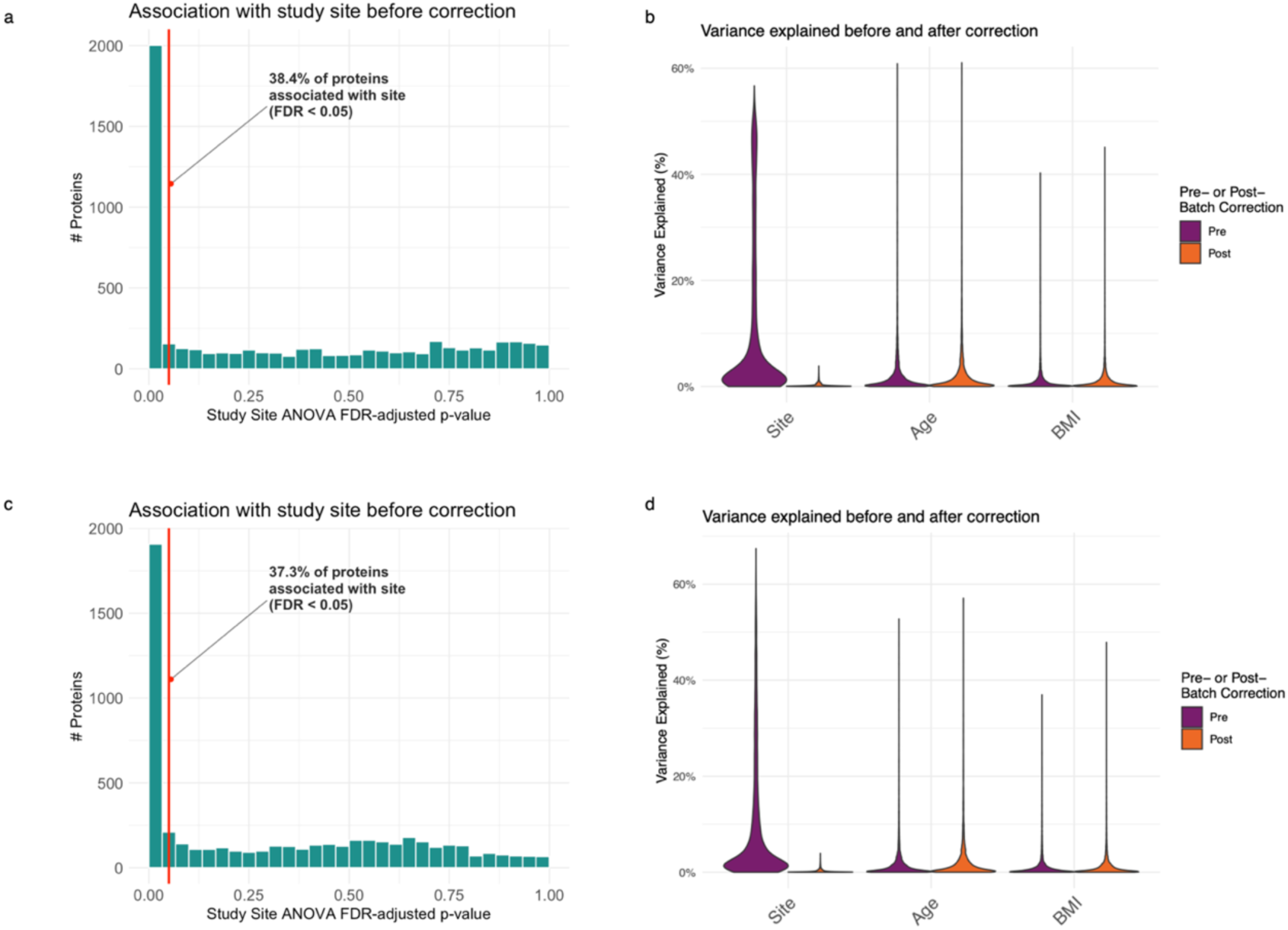
Batch correction preserves biological variance: **a**, variancePartition variance decomposition before ComBat on the normalized matrix. Distribution of ANOVA FDR-adjusted p-values for the association between each protein and study site. The red vertical line indicates the 0.05 FDR significance threshold. Prior to correction, 38.4% of proteins showed a significant association with study site (ANOVA, BH-adjusted P < 0.05). **b,** Variance explained by covariates before and after batch correction. Violin plots show the percentage of variance in protein abundance explained by each covariate (Site, Age, BMI) before (“Pre”) and after (“Post”) batch correction. Batch correction effectively removes site-driven variance while preserving variance attributable to biological factors such as age and BMI. **c,** variancePartition variance decomposition before and after ComBat on the EVA input matrix. Distribution of ANOVA FDR-adjusted p-values for the association between each protein and study site. The red vertical line indicates the 0.05 FDR significance threshold. Prior to correction, 37.3% of proteins showed a significant association with study site (ANOVA, BH-adjusted P < 0.05). **d,** Variance explained by covariates before and after batch correction. Violin plots show the percentage of variance in protein abundance explained by each covariate (Site, Age, BMI) before (“Pre”) and after (“Post”) batch correction. Batch correction effectively removes site-driven variance while preserving variance attributable to biological factors such as age and BMI.

**Supplementary Figure 2:**
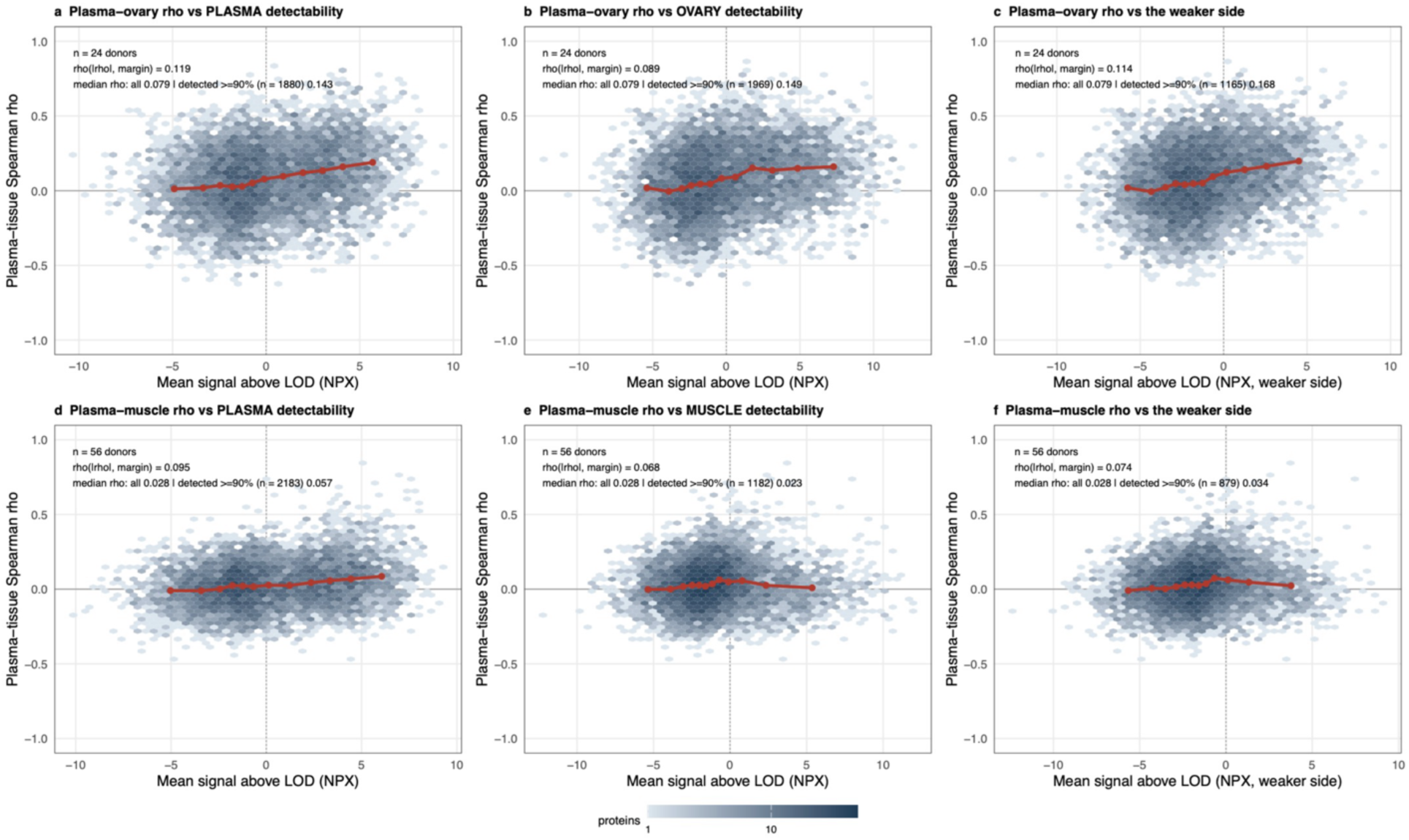
Plasma–tissue correlation against detectability relative to the fixed limit of detection. Each panel plots the covariate-adjusted plasma–tissue Spearman ρ of a protein (y) against its mean signal above the Olink fixed LOD (x, NPX units), as a hexagonal-binned density; the red line joins the medians of twelve equal-count bins of the x axis. Detectability is computed over exactly the participants that entered the correlation. **a–c,** plasma–ovary (24 participants): ρ against detectability on the plasma side **a**, on the ovary side **b**, and against the weaker of the two sides (**c**, the per-protein minimum, shown as a summary rather than as a compartment). **d–f,** plasma–muscle (56 participants), same three views. Annotations give the Spearman correlation between |ρ| and mean signal above LOD, and the median ρ of all 5,415 proteins versus of the well-detected subset (detection rate ≥ 90%). The relationship is positive but shallow in every panel (ρ(|ρ|, margin) = 0.119, 0.089 and 0.114 for ovary; 0.095, 0.068 and 0.074 for muscle), and restricting to well-detected proteins raises the median ρ only from 0.079 to 0.143 (ovary, n = 1,880) and from 0.028 to 0.057 (muscle, n = 2,183). Measurement sensitivity therefore accounts for part of the spread but not for the near-zero central tendency.

**Supplementary Figure 3:**
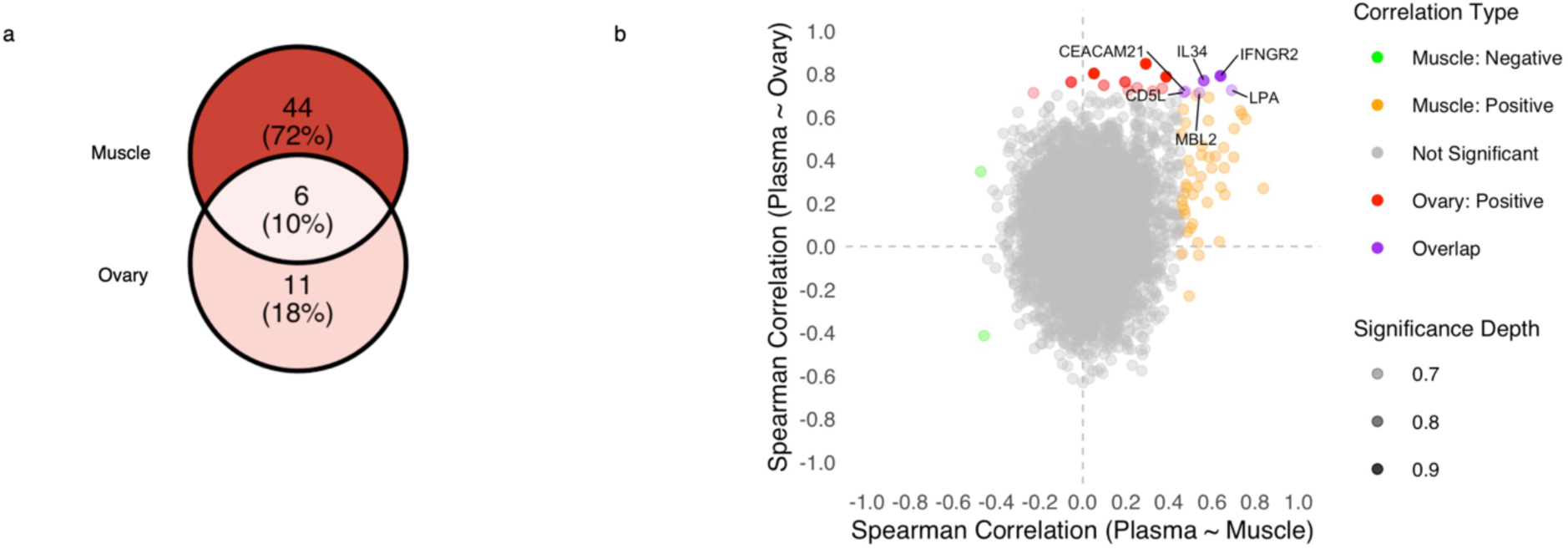
Proteins overlap in plasma-ovary correlation and plasma-muscle correlation: **a**, Distribution of significant plasma–tissue protein correlations across ovary and muscle. Green dots represent significant negative correlations in plasma–muscle, orange dots represent significant positive correlations in plasma–muscle, red dots represent significant positive correlations in plasma–ovary, purple dots indicate proteins significantly correlated in both plasma–muscle and plasma–ovary, and grey dots indicate non-significant correlations. **b,** Overlap of significant protein correlations between plasma–muscle and plasma–ovary, highlighting six proteins shared between the two comparisons.

**Supplementary Figure 4:**
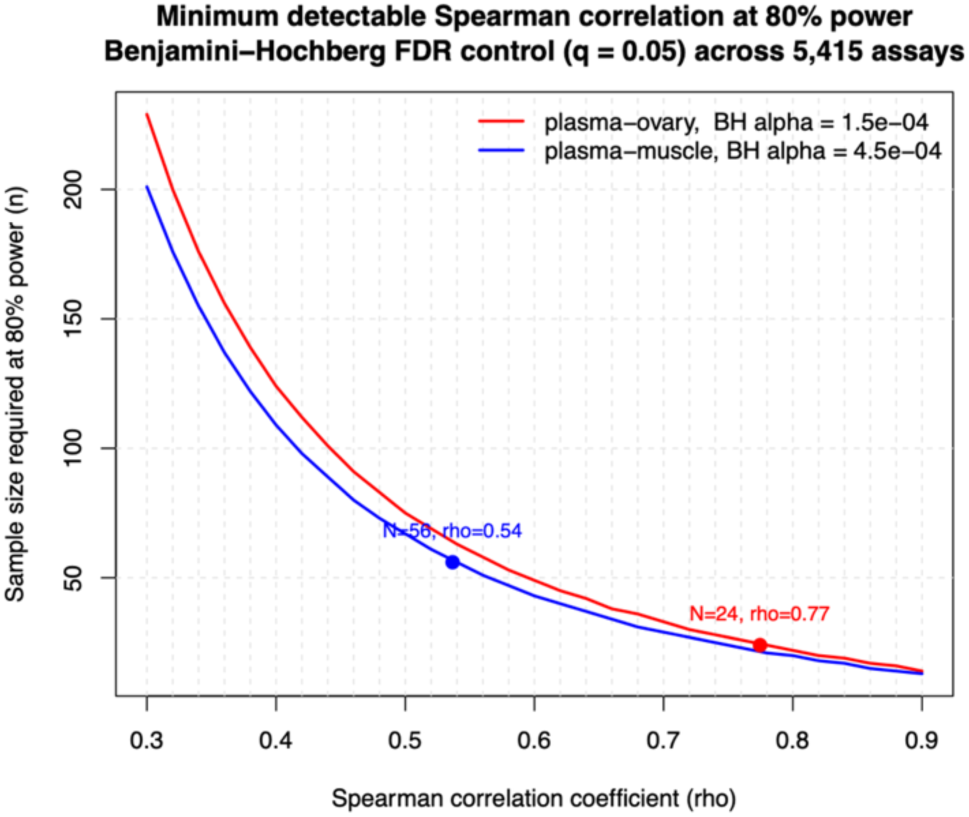
Power analysis for spearman correlation. Sample size required to detect a plasma–tissue Spearman correlation at 80% power, computed with pwr.r.test under a two-sided test. The significance level is the one the analysis actually enforces — the largest raw P still called significant by the Benjamini–Hochberg procedure at q < 0.05 within each arm (1.5 × 10⁻⁴ for plasma–ovary, 4.5 × 10⁻⁴ for plasma–muscle). Curves give the required N across ρ; annotated points mark the two matched cohorts. At their realised sizes the ovary arm (N = 24) is powered to detect |ρ| ≥ 0.77 and the muscle arm (N = 56) |ρ| ≥ 0.54; the smallest coefficients that in fact passed BH correction were |ρ| = 0.71 and 0.46 respectively.

**Supplementary Figure 5:**
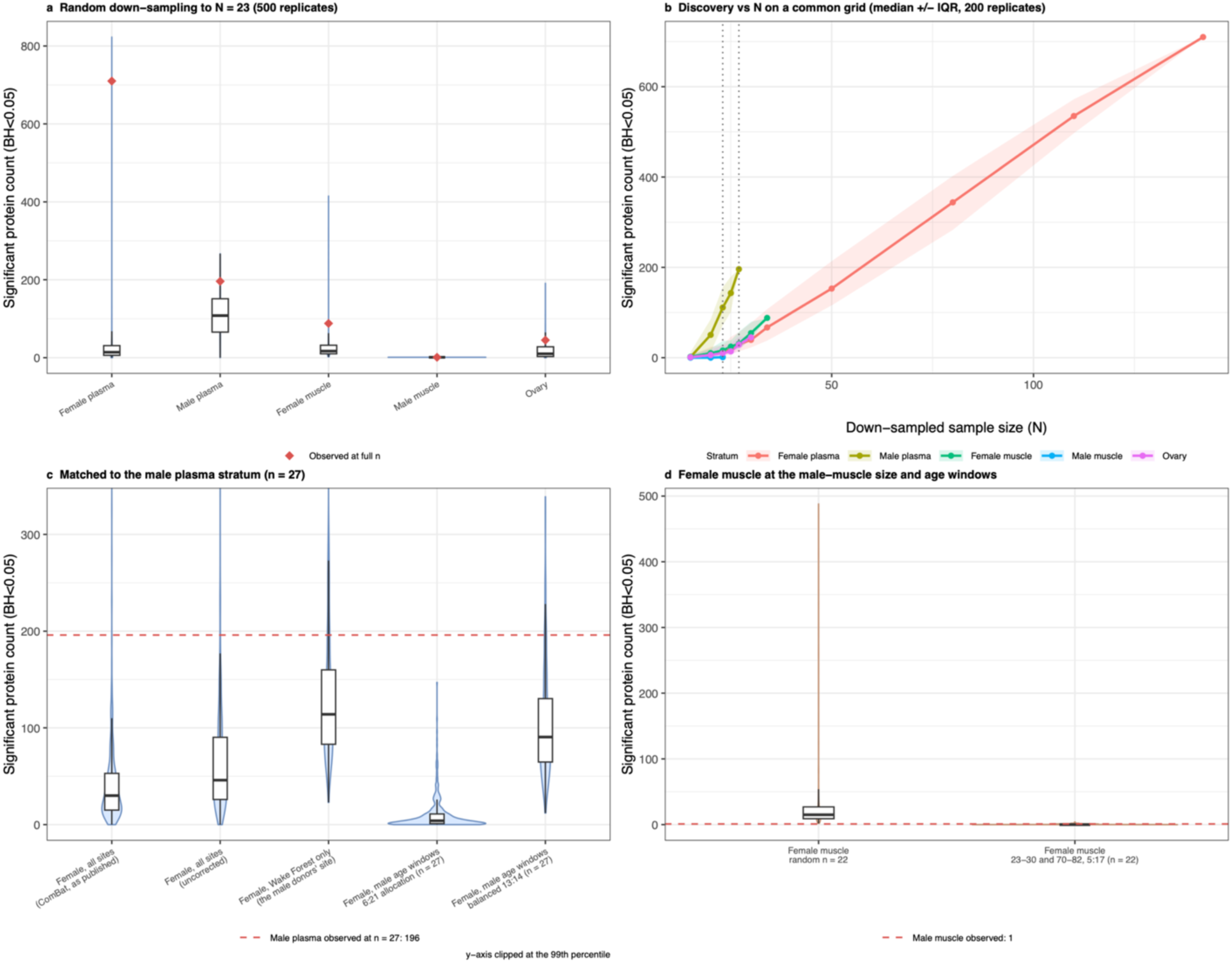
Sensitivity analyses for robustness of age-correlated protein discovery: **a**, Down-sampling each stratum to N=23 across 500 replicates shows that discovery rank order is not driven by unequal sample sizes. **b,** Down-sampling curves (Spearman age-association, BH<0.05) for each stratum; line = median, ribbon = IQR over 200 replicates, black diamond = full-cohort value. Female plasma continues to rise without plateau, indicating discovery is sample-size-limited; the smaller cohorts are bounded by their available N. **c,** Female plasma resampled to the male plasma stratum’s own n = 27 under five designs, against the male observed count of 196 (dashed line): all sites with ComBat as published (median 30), all sites uncorrected (46), Wake Forest University only, the site all male participants come from (114), the male age windows at the male 6 : 21 allocation (4), and the same windows at a balanced 13 : 14 allocation (90.5). **d,** Female muscle at the male muscle size and age windows (23–30 and 70–82 yr, 5 : 17, n = 22; median 0) against a random draw of the same size (median 15) and the male muscle observed count of 1. Violins show the resampling distribution, boxes the interquartile range; 500 replicates per design.

**Supplementary Figure 6:**
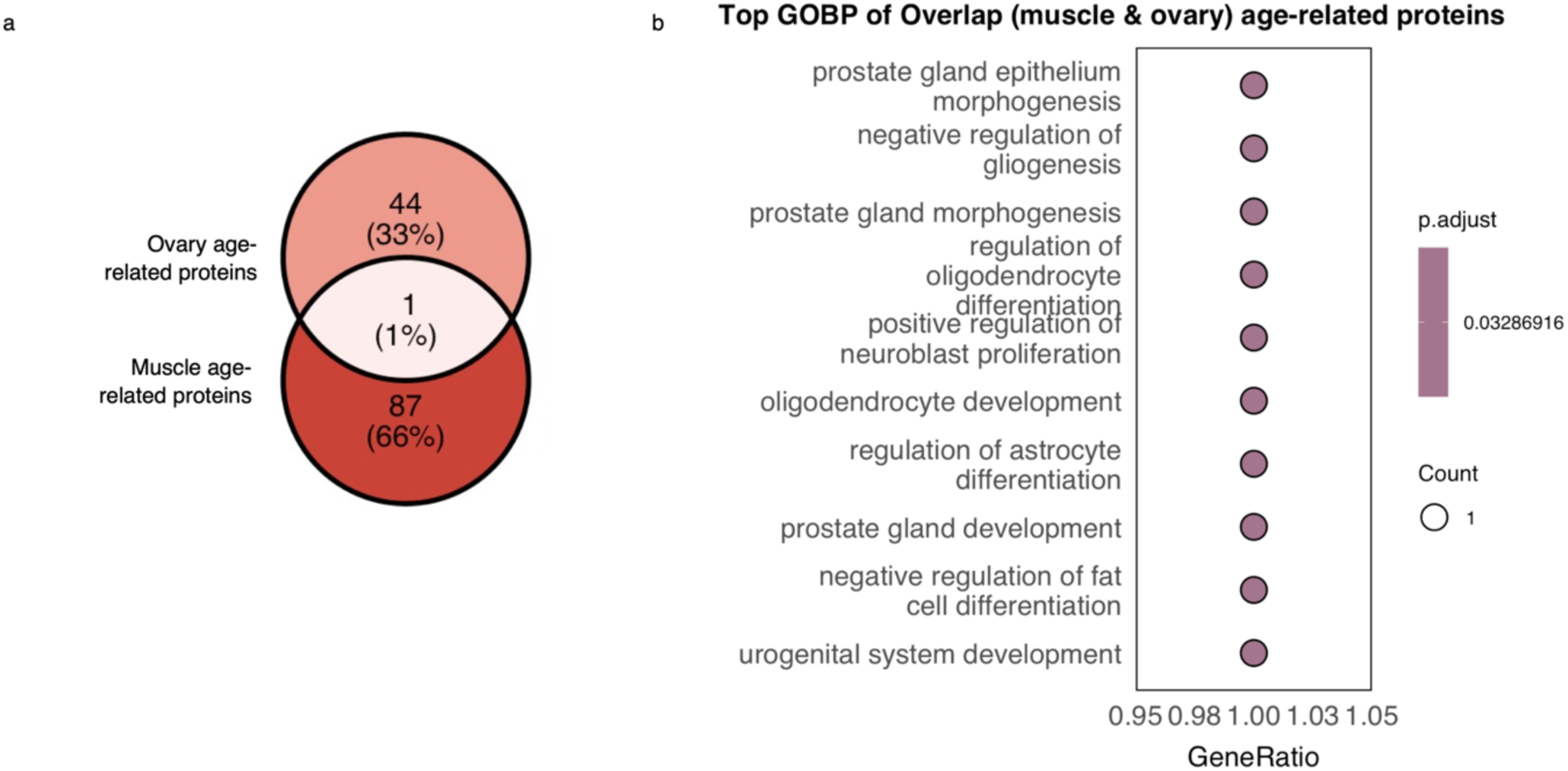
Tissue-specific proteomic signatures of aging. **a**, Overlap of age-correlated proteins between ovary and muscle (union of female and male muscle): 44 proteins were ovary-specific, 87 muscle-specific and only 1 shared, underscoring that age-correlated proteins are largely tissue-private. **b,** Gene Ontology biological processes enriched (over-representation analysis, hypergeometric test with Benjamini–Hochberg correction) among the age-correlated proteins shared between muscle and ovary.

**Supplementary Figure 7:**
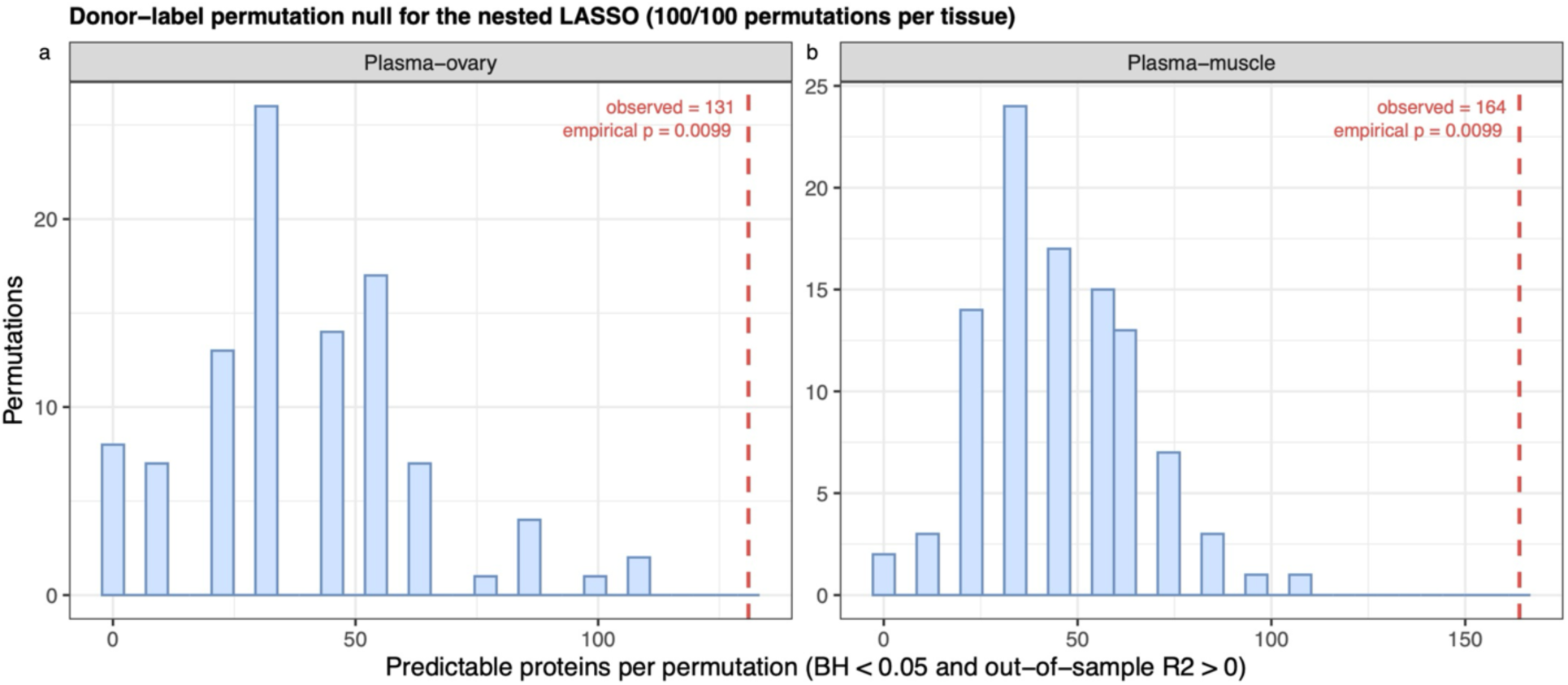
Donor-label permutation null for the nested plasma-to-tissue LASSO. Distribution of the number of predictable proteins (raw P at or below the BH cutoff realized in the observed run and out-of-sample R² > 0) obtained in 100 permutations per tissue, in which tissue donor labels were shuffled before covariate residualization which also breaks the tissue–covariate correspondence. Each permutation used one 5-fold outer cross-validation over a fixed abundance-stratified panel of 500 targets, and counts were scaled to the full 5,415-protein panel. Dashed line, the observed count. **a,** Plasma–ovary: 131 observed against a null mean of 39.2 (s.d. 23.6, maximum 108.3). **b,** Plasma–muscle: 164 against 45.1 (s.d. 21.1, maximum 108.3). Empirical P = 0.0099 in both tissues, the smallest value attainable with 100 permutations.

**Supplementary Figure 8:**
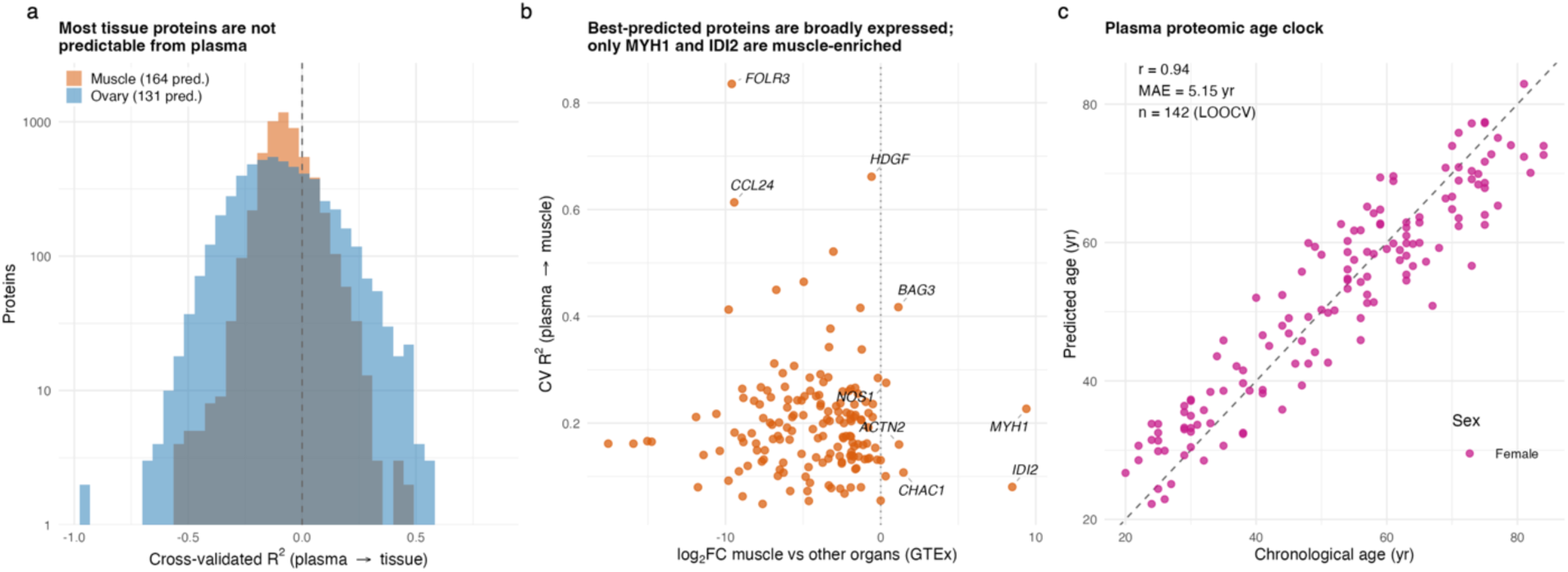
Chronological age. **a**, Distribution of cross-validated R² (plasma→tissue) across all tested proteins for muscle (164 predictable) and ovary (131 predictable); most proteins are not predictable from plasma. **b,** GTEx tissue-of-origin versus predictability for muscle: cross-validated R² (plasma→muscle) against log₂ fold-change of muscle-vs-other-organ expression. The best-predicted proteins (FOLR3, CCL24, HDGF) are secreted or broadly expressed rather than muscle-private, whereas bona fide muscle-specific proteins (MYH1, MYL1, ENO3, IDI2, SMTNL2, BIN1) fall at high positive log₂FC. **c,** Exploratory plasma age model: fitted versus chronological age under leave-one-out cross-validation (female, ComBat-adjusted; n = 142; r = 0.94, MAE ≈ 5.15 yr). Cohort-wide age-preserving ComBat correction preceded cross-validation, so these statistics do not constitute independent validation. Covariate residualization and imputation are fitted inside each training fold; the counts are shown against a 100-permutation donor-label null.

**Supplementary Figure 9:**
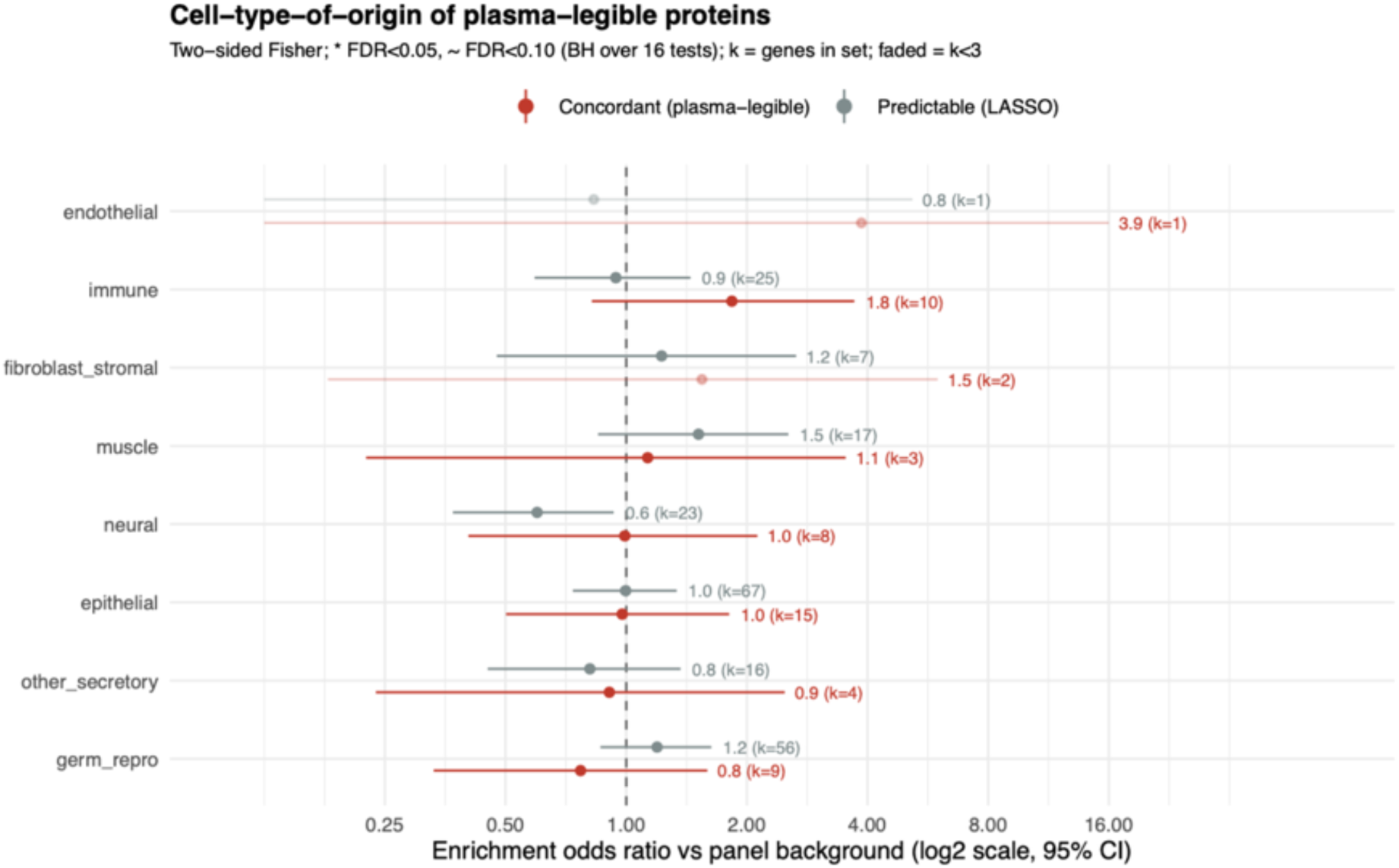
Cell-type-of-origin of plasma-legible proteins (Human Protein Atlas single-cell atlas). Each analyte was assigned the cell lineage of its most cell-type-specific single-cell expression in HPA (immune, endothelial, epithelial, muscle, fibroblast/stromal, neural, germ/reproductive, other secretory). The forest plot shows the enrichment odds ratio (two-sided Fisher’s exact test; log2 scale, 95% CI) of each lineage among plasma–tissue concordant proteins (red) and the broader LASSO-predictable set (grey) versus the annotated panel background; k = number of set genes in the lineage, points with k < 3 are faded, and significance is BH-adjusted across all 16 tests (* FDR < 0.05, ∼ FDR < 0.10). Concordant proteins show a nominal immune skew (OR = 1.8, k = 10) that does not survive multiple-testing correction, whereas the predictable set sits at background across lineages; this analysis is exploratory given the small concordant set (n = 57).

**Supplementary Figure 10.**
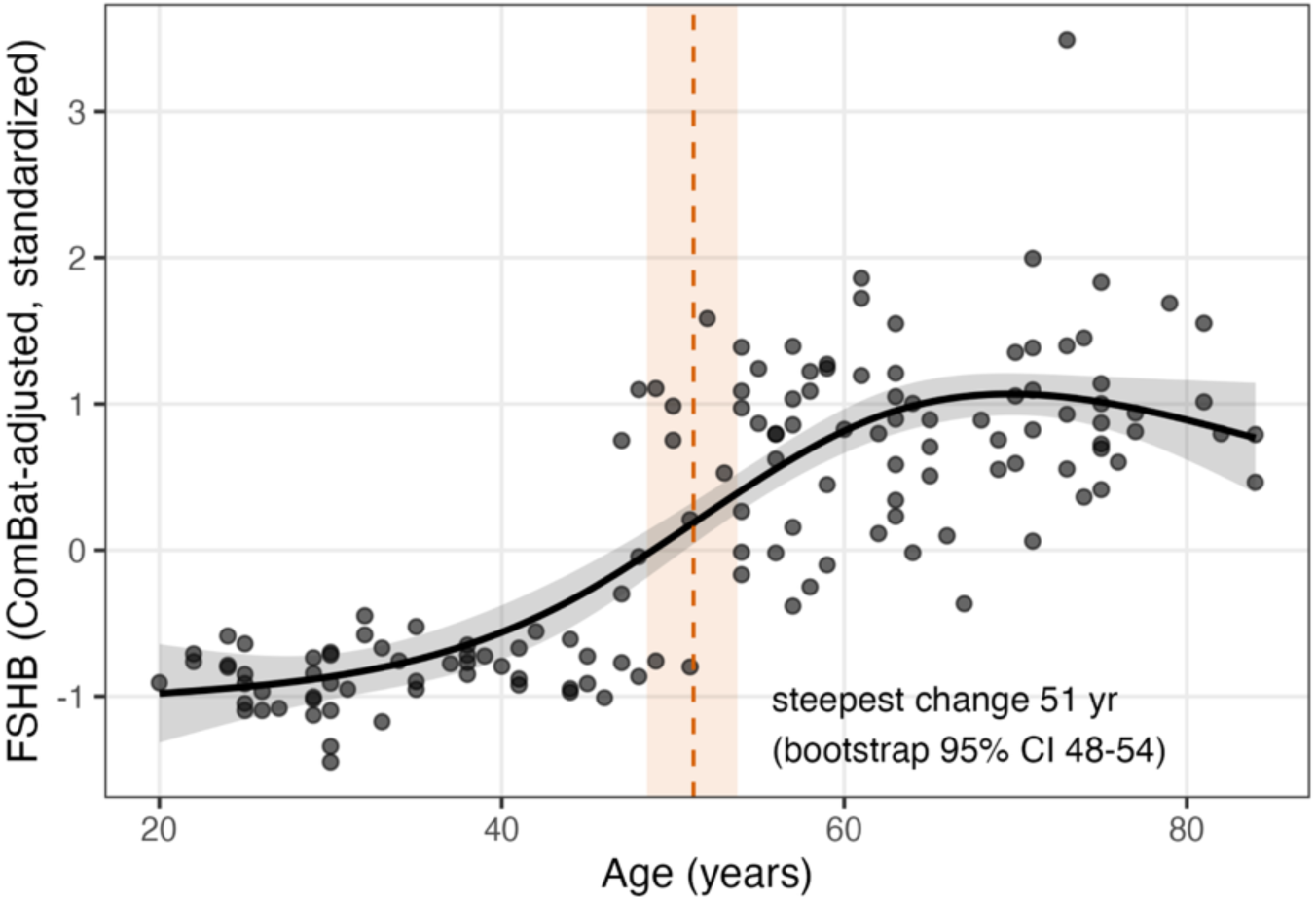
Plasma FSHB rises sharply across the menopausal transition in females. Among eight reproductive-axis candidate proteins screened in female plasma (FSHB, LHB, SHBG, PRL, INHBB, INHBC, WFDC2, MUC16), FSHB was the only protein with a well-defined age of maximal change and is shown here. Expression is ComBat-adjusted, standardized Olink protein level; each grey point is one female plasma sample. The solid black line is a natural cubic-spline fit of expression on age (natural spline, 3 df) adjusted for BMI, evaluated at the median BMI, and the grey band is its pointwise 95% confidence interval. The vertical orange dashed line marks the age of steepest change (the maximum absolute first derivative of the fitted spline), and the orange shaded band is the corresponding bootstrap 95% confidence interval (2,000 resamples). FSHB rises sigmoidally, with steepest change at 51 years (bootstrap 95% CI 48–54), consistent with the menopausal FSH surge.

**Supplementary Figure 11:**
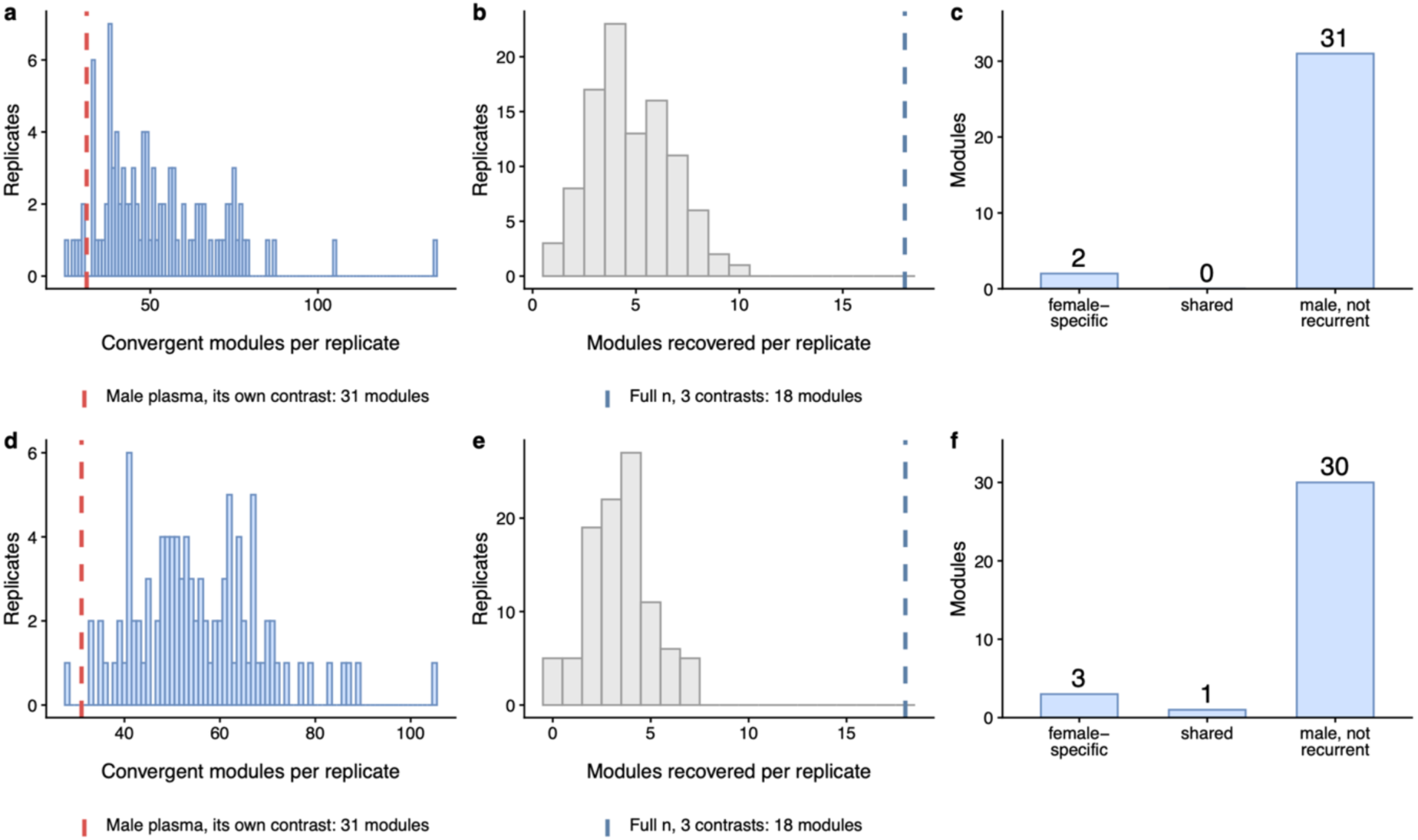
Female plasma matched to the male plasma design, with and without the site and ComBat confound. **a**, Number of convergent modules per replicate when female plasma is down-sampled 100 times to a single young-versus-old contrast matched to the male contrast on allocation (6 young : 21 old, n = 27) and on age window (young drawn from 23–30 and old from 70–87 years, the ranges the male donors span). Dashed line, male plasma observed at its own n. The fraction of replicates at or below the male count is an empirical tail proportion, not a P value: the male value is a single observation, not a draw from this distribution. **b,** How many of the female convergent modules reported at full n are re-identified per replicate; the reference (dashed line) is the union of the three female contrasts at full n. A down-sampled replicate runs one contrast, so modules arising only from the other two contrasts cannot be recovered by it, and the count against the single young-versus-old set is carried alongside. **c,** Module identity of the down-sampled female set relative to the male set: female-specific, recurrent in female and absent from the male set; shared, recurrent in female and present in the male set; male, not recurrent, in the male set but not recurrent in female. A module counts as recurrent when it is significant in at least 50% of the replicates that tested it. **d–f,** as a–c, but using female plasma from Wake Forest University only and without ComBat correction; all 27 male donors come from this site, so this arm removes the site and batch-correction difference between the sexes.

**Supplementary Fig. 12:**
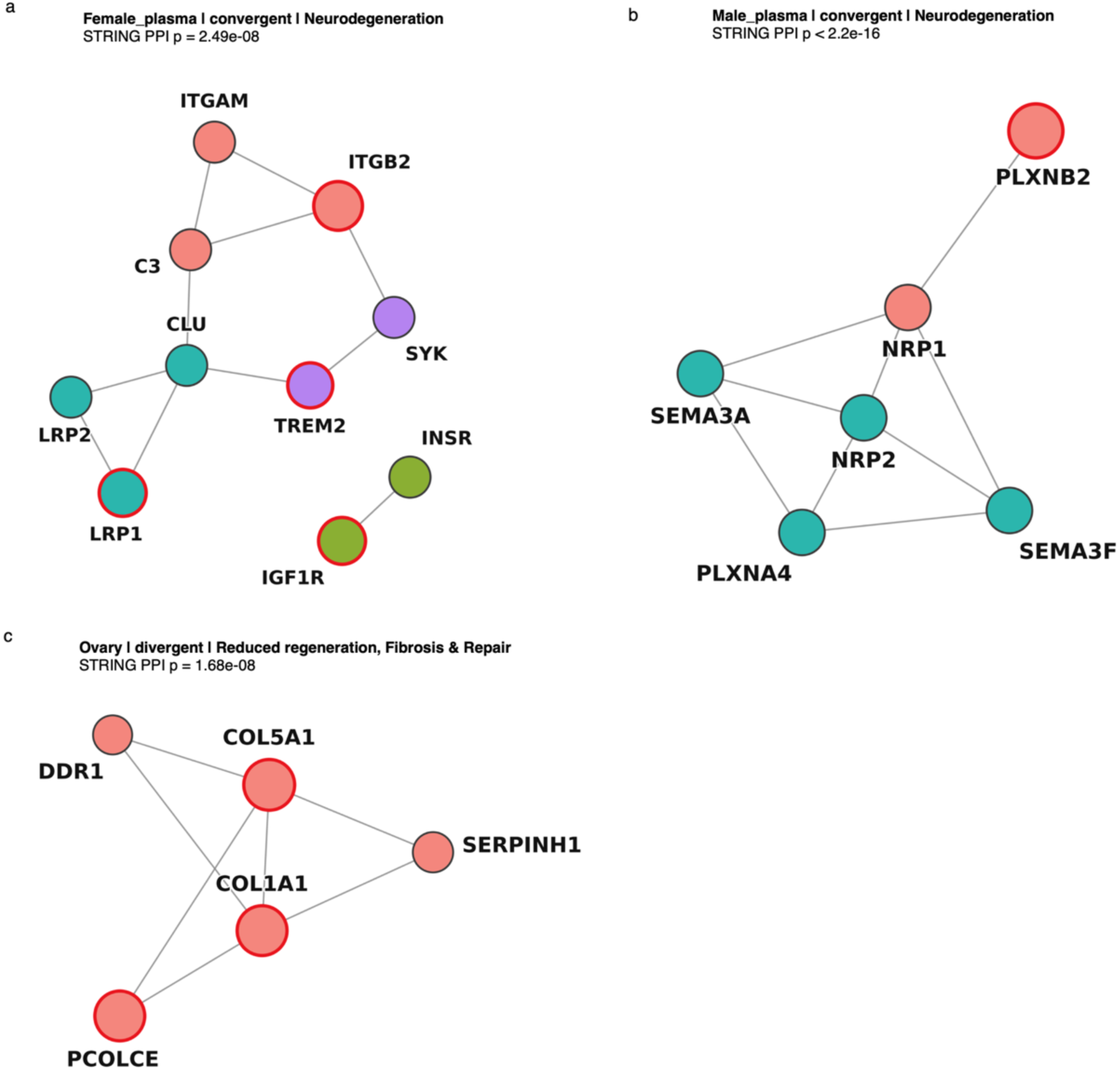
Protein–protein interaction (PPI) networks of selected age-reordered functional modules. **a**, PPI network of the age-convergent Neurodegeneration module in female plasma. **b,** PPI network of the age-convergent Neurodegeneration module in male plasma. **c,** PPI network of the age-divergent Reduced Regeneration, Fibrosis & Repair module in ovary. Nodes represent proteins and edges represent STRING protein–protein associations.

